# Identification of highly cross-reactive mimotopes for a public T cell response in murine melanoma

**DOI:** 10.1101/2022.02.28.482028

**Authors:** Beth E. Grace, Coralie M. Backlund, Duncan M. Morgan, Byong H. Kang, Nishant K. Singh, Brooke D. Huisman, C. Garrett Rappazzo, Kelly D. Moynihan, Laura Maiorino, Connor S. Dobson, Taeyoon Kyung, Khloe S. Gordon, Patrick V. Holec, Overbeck C. Takou Mbah, Daniel Garafola, Shengwei Wu, J. Christopher Love, K. Dane Wittrup, Darrell J. Irvine, Michael E. Birnbaum

## Abstract

While immune checkpoint blockade results in durable responses for some patients, many others have not experienced such benefits. These treatments rely upon reinvigorating specific T cell-antigen interactions. However, it is often unknown what antigens are being recognized by T cells or how to potently induce antigen-specific responses in a broadly applicable manner. Here, we characterized the CD8^+^ T cell response to a murine model of melanoma following combination immunotherapy to determine the basis of tumor recognition. Sequencing of tumor-infiltrating T cells revealed a repertoire of highly homologous TCR sequences that were particularly expanded in treated mice and which recognized an antigen from an endogenous retrovirus. While vaccination against this peptide failed to raise a protective T cell response *in vivo*, engineered antigen mimotopes induced a significant expansion of CD8^+^ T cells cross-reactive to the original antigen. Vaccination with mimotopes resulted in killing of antigen-loaded cells *in vivo* yet showed modest survival benefit in a prophylactic vaccine paradigm. Together, this work demonstrates the identification of a dominant tumor-associated antigen and generation of mimotopes which can induce robust functional T cell responses that are cross-reactive to the endogenous antigen across multiple individuals.

## Introduction

Immune checkpoint blockade (ICB) therapies such as anti-PD-1 and anti-CTLA-4 antibodies have revolutionized the field of immunotherapy. These therapies work by reinvigorating pre-existing anti-tumor T cell responses in cancer patients (1). However, it is often difficult to identify the antigen or set of antigens that is being targeted in ICB therapies. While patient-specific neoantigens represent a significant proportion of anti-tumor immune responses (2), there are also shared antigens that can each contribute to strong anti-tumor immunity, including common oncogenic mutations, tissue lineage antigens, and viral antigens (3,4). Thus, identification and characterization of antigens that correlate with successful ICB therapies is essential. Recognition of these antigens may represent a prognostic marker for the potential success of ICB therapies, or the antigens may themselves be targeted in combination therapies with the aim of further expanding the pool of patients that benefit from the extended survival these therapies can offer (5).

The combination of antigen-targeted vaccines and ICB is expected to be synergistic, since vaccines can expand tumor-specific T cells and ICB can allow the T cell response to proceed uninhibited by checkpoint mediated suppression (6). Many classes of putative tumor antigens tend to activate their cognate T cells less potently than pathogen-derived antigens, at least in part due to deletion of the most potent T cell clones against self-antigens via thymic selection (7). Peptide mimics of an antigen, referred to as mimotopes or altered peptide ligands, may be able to induce a stronger T cell response to the native antigen by more potently engaging with a given T cell receptor (TCR) than the native antigen (8,9). Given the extensive cross reactivity of T cells (10–12), mimotopes can be used to expand a population of T cells that overlap with the population that recognize the native antigen (13,14). In this way, mimotopes can be utilized as a vaccine, resulting in the expansion of a population of reactive T cells that can also recognize the native antigen when it is presented by tumor cells. Mimotopes have also previously been used to mobilize low-avidity self-specific T cells (8,15).

Here, we investigate the tumor-infiltrating T cell response following combination therapy in the well-studied, poorly immunogenic B16F10 murine model of melanoma. We identify a set of strikingly similar T cell clones shared both within individual immune repertoires and between animals and determine the antigen recognized by these clones. We then use a yeast display pMHC library approach to identify and characterize two antigen mimotopes that potently activate T cells. This work supports the utility of mimotopes for the improvement and expansion of T cell responses against tumor-associated antigens.

## Results

### The immune response following combination immunotherapy is oligoclonal and includes a set of highly conserved T cell clones

We first set out to characterize the T cell response in C57BL/6 mice following treatment of B16F10 melanoma with a combination therapy that has been previously shown to induce a robust protective immune response. The therapy, known as AIP, combines the Trp1-targeting antibody TA99 (A), extended half-life IL-2 (I), and an anti-PD-1 antibody (P), and results in cures in about half of mice with B16F10 melanoma (16). While a four-component version of the therapy, known as AIPV, also includes an amphiphile peptide vaccine targeting Trp2, we focused upon AIP in this study as it is more representative of current combination ICB therapies and does not direct the response to a particular antigen. Further, previous studies demonstrated that in the absence of a peptide vaccine component, AIP is effective yet primarily targets an antigen that is distinct from Trp2 (16). We single-cell sorted and sequenced the T cell receptors of activated tumor-infiltrating lymphocytes (TILs) in either AIP-treated or untreated mice. The AIP therapy was administered on days 8 and 15 following tumor inoculation, and IFN-γ-secreting CD8^+^ TILs were sorted on day 21 and sequenced **(Figure 1A).**

**Figure 1:**
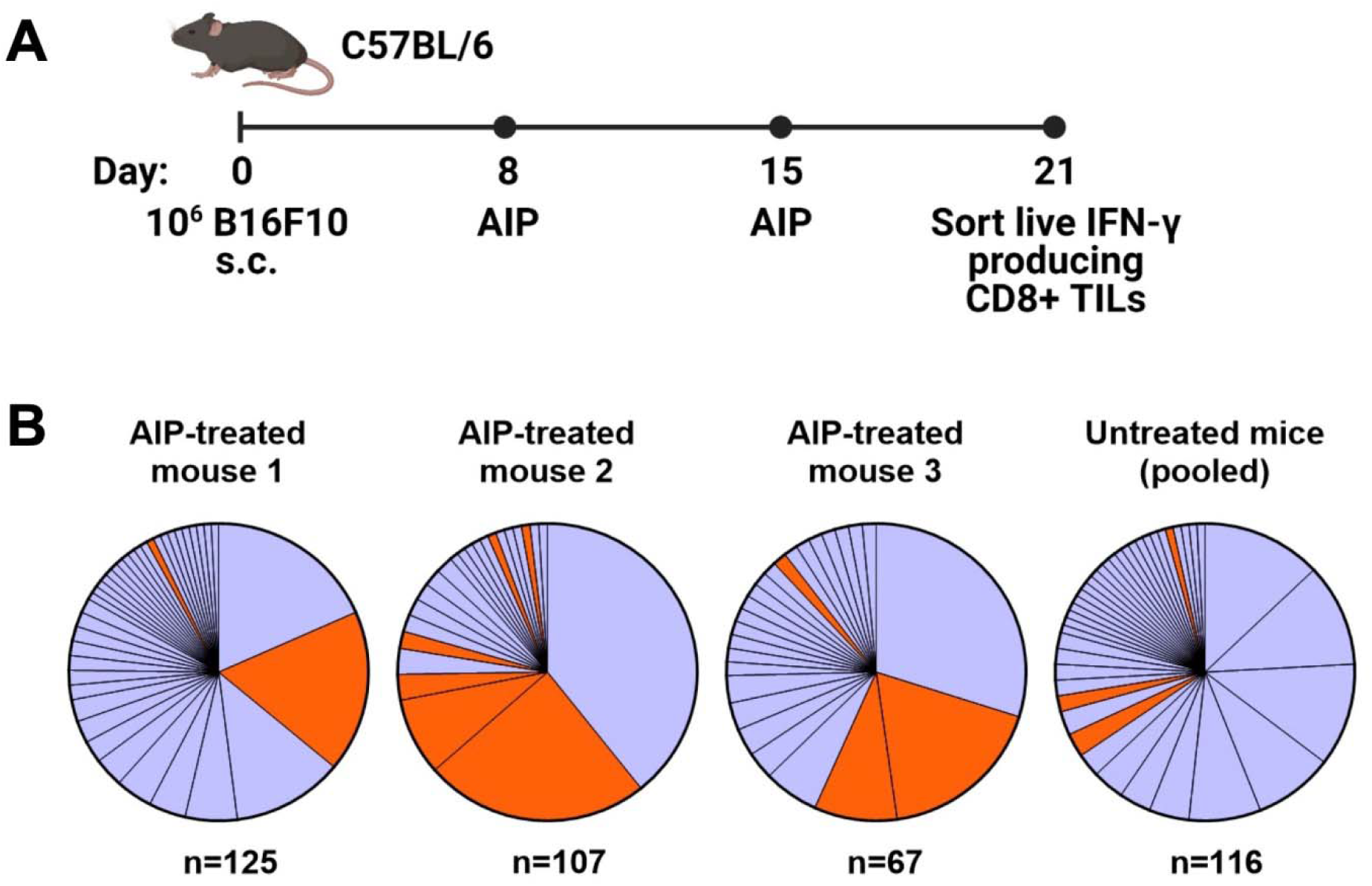
The T cell response following combination immunotherapy of melanoma is oligoclonal and contains a set of homologous T cell clones. (A) Timeline of tumor inoculation, therapy, and cell sorting (n=5 mice/group, s.c. = subcutaneous). (B) Clonality of T cell responses in individual treated mice and pooled untreated mice. T cells with identical alpha and beta chain sequences were identified as members of a clone. N indicates the number of T cells for which paired TCR sequences were obtained. Orange colored clones appear in Table 1.

Analysis of the TCR repertoire from recovered TILs demonstrated an oligoclonal immune response, with 40-75% of the IFN-γ^+^ CD8^+^ T cell response consisting of three to four dominant clones. Interestingly, both AIP-treated and untreated mice demonstrated similarly oligoclonal responses **(Figure 1B).** The presence of multiple large clones may indicate that the untreated mice are mounting an immune response to the tumor, albeit not one effective enough to reduce tumor burden without additional intervention. In AIP-treated mice, CD8^+^ T cell infiltration in tumors was tenfold higher than in untreated mice **(Supplementary Figure 1A),** suggestive of highly expanded T cell clones in AIP-treated mice, although the fraction of CD8^+^ T cells that was activated was similar in treated and untreated mice **(Supplementary Figure 1B).**

Upon analysis of CDR3 sequences of the T cells, we identified two highly similar CDR3α sequences which were highly represented in our data **(Table** 1); clones containing these sequences are colored orange **(Figure 1B).** Each of these sequences relied upon TRAJ7 in forming the CDR3 sequence, with a small degree of variation due to V-J junctional diversity. These CDR3α sequences were present in several of the dominant clones, particularly in treated mice, while also appearing occasionally in the untreated pool. In addition to the strikingly similar CDR3 regions, the expanded TCRs showed a restricted TRAV repertoire, with the variable regions having similarities in their CDR1 and CDR2 sequences **(Supplementary Figure 1C).**

**Table 1:** CDR3 sequences and gene usage for T cells with strikingly similar TCR sequences. The clone size and mouse in which the clone was identified are also shown.

| Clone Name | Clone Size | Mouse | CDR3α | TRAV | TRAJ | CDR3β | TRBV | TRBD | TRBJ |
| --- | --- | --- | --- | --- | --- | --- | --- | --- | --- |
| 7PPG1 | 26 | AIP 2 | CAAKDYSNNRLTL | 7-2 | 7 | CASS**PP**GD**T**QYF | 3 | 1 | 2-5 |
| 7PPG2 | 22 | AIP 1 | CAAKDYSNNRLTL | 10 | 7 | CASS**PP**GGSEVFF | 5 | 1 | 1-1 |
| 7PPG3 | 12 | AIP 3 | CAAKDYSNNRLTL | 10 | 7 | CASS**PP**GSQNTLYF | 5 | 1 | 2-4 |
| 7ELG1 | 9 | AIP 2 | CAASDYSNNRLTL | 7-2 | 7 | CASS**L****EL**GGREQYF | 16 | 2 | 2-7 |
| 7PPG4 | 6 | AIP 3 | CAAKDYSNNRLTL | 10D | 7 | CASS**PP**GQNT**E**VFF | 4 | 1 | 1-1 |
| 7DLG1 | 3 | untr | CAAKDYSNNRLTL | 10D | 7 | CASSQ**DL**GN**S**DYTF | 5 | 1 | 1-2 |
| 7DLG2 | 3 | AIP 2 | CAAKDYSNNRLTL | 10D | 7 | CASSQ**DL**GY**A**EQFF | 5 | 2 | 2-1 |
| 7AQG1 | 2 | untr | CAASDYSNNRLTL | 7-2 | 7 | CASSQ**A**QGGG**A**EQFF | 5 | 1 | 2-1 |
| 7EGA1 | 2 | AIP 2 | CAAKDYSNNRLTL | 5D-4 | 7 | CASSQ**E**G**A**NTEVFF | 2 | 1 | 1-1 |
| 7PPG5 | 1 | untr | CAAKDYSNNRLTL | 10 | 7 | CASS**PP**GVNTEVFF | 5 | 1 | 1-1 |
| 7DPG1 | 1 | AIP 1 | CAAKDYSNNRLTL | 10 | 7 | CASSQ**D**PGGN**A**EQFF | 2 | 2 | 2-1 |
| 7PPG6 | 1 | AIP 3 | CAAKDYSNNRLTL | 10 | 7 | CASS**PP**GGDTEVFF | 5 | 1 | 1-1 |
| 7TPG1 | 1 | AIP 2 | CAAKDYSNNRLTL | 7-2 | 7 | CAS**S****T**PGQNT**E**VFF | 4 | 1 | 1-1 |
| 7NWS1 | 1 | AIP 2 | CAASDYSNNRLTL | 7-2 | 7 | CASS**L****N**WSQDTQYF | 16 | 2 | 2-5 |

The beta chains paired with these alpha chains varied overall, but there were some similarities among the CDR3β sequences, such as a PPG motif present in 7 out of 14 sequences and frequent usage of TRBV5. Based on the resemblances among these clones overall, we hypothesized that they recognize a shared antigen. Since they were also particularly expanded in treated mice, we chose to focus on this set of clones for antigen identification.

### Prominent T cell clones recognize an endogenous retrovirus peptide presented by multiple tumor types

To identify the cognate antigen of these TILs, we first tested whether cells expressing the identified TCRs selectively recognized tumor cell lines and if that recognition correlated with expression of melanocyte tissue lineage antigens. Murine CD8^+^ 58^-^/^-^ T cell hybridomas were transduced with the 7PPG2 and 7PPG4 TCRs, two representative TIL sequences sharing TCR alpha chain homology but expressing distinct TCR beta chains. The transduced T cell lines were then tested for activation as measured by IL-2 secretion upon challenge by a panel of tumor cell lines pre-treated with IFN-γ to upregulate class I MHC expression **(Figure 2A, Supplementary Figure 2A).** As a positive control, 58^-^/^-^ T cells transduced with the 2C TCR were challenged via coculture assay with B16.SIY cells, a B16F10 melanoma line modified to express the 2C cognate antigen SIY (17). We found that 7PPG2- and 7PPG4-transduced 58^-^/^-^ T cells were activated following co-culture with B16F10 and B16.SIY cells but not TC-1 cells, a transformed mouse lung epithelial cell line **(Figure 2A).** Intriguingly, 58^-^/^-^ T cells transduced with 7PPG2 and 7PPG4, but not 2C, also recognized MC-38 cells, a murine colon adenocarcinoma cell line **(Figure 2A).** Therefore, 7PPG2 and 7PPG4 TCR-transduced 58^-^/^-^ T cells showed a degree of antigen selectivity that did not appear in only melanocyte-specific contexts.

**Figure 2:**
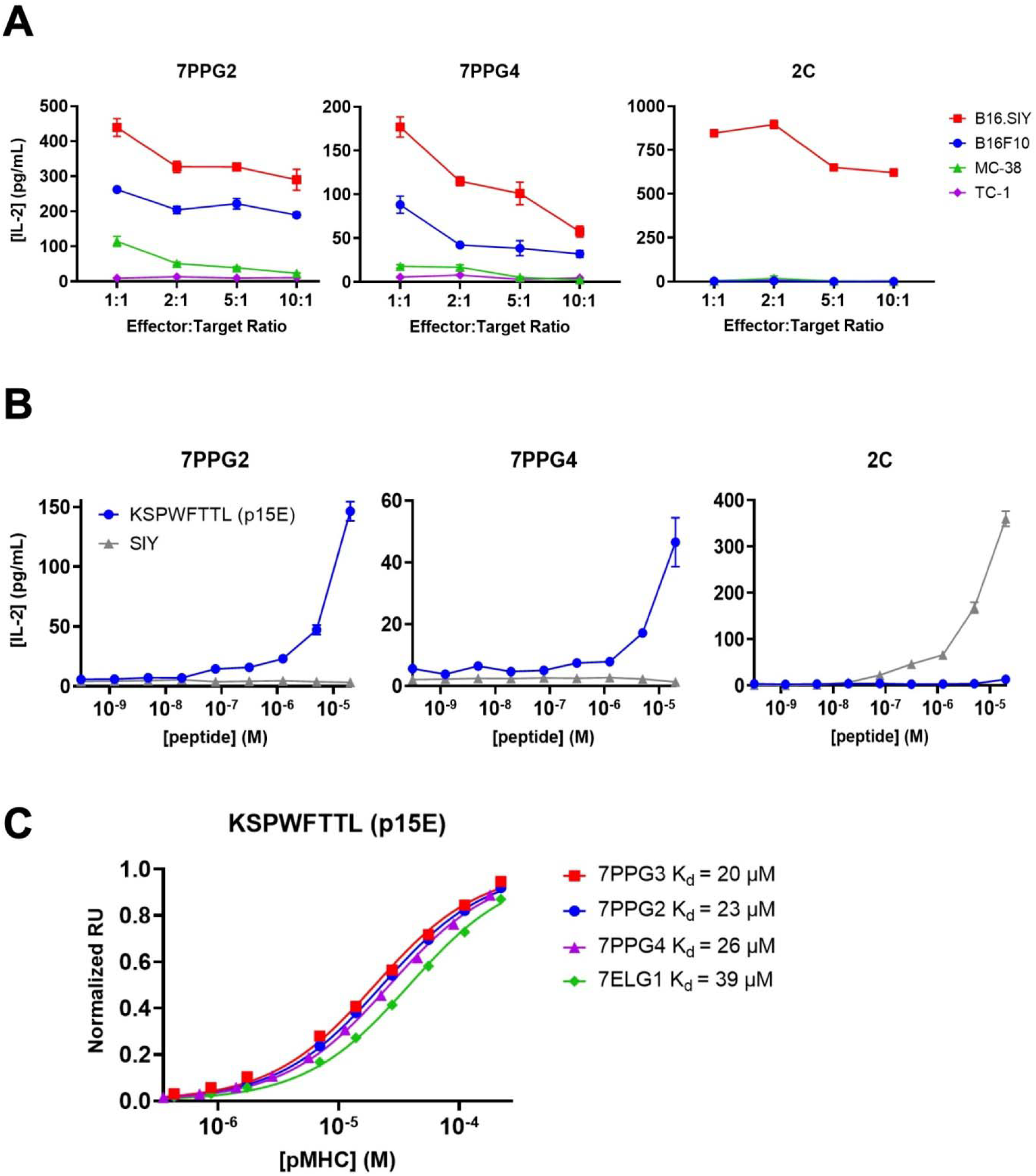
TCR-transduced T cells are activated by an endogenous retrovirus peptide expressed by multiple tumor types. (A-B) 58^-/-^ cells transduced to express the 7PPG2, 7PPG4, and 2C TCRs were cocultured with B16F10, B16.SIY, MC-38, and TC-1 cancer cell lines at various effector:target ratios (A) or with DC2.4 cells loaded with p15E or SIY peptide (B). T cell activation was assessed by IL-2 ELISA. Data shown are mean+s.e.m. of triplicate (A) or duplicate (B) samples and are representative of 3 independent experiments. (C) Surface plasmon resonance was performed to quantify affinities between four TCRs and KSPWFTTL peptide presented by H-2K^b^.

A recent study examining the humoral immune responses in AIPV-treated mice noted that the majority of the induced antibody response targeted an endogenous retroviral envelope glycoprotein from murine leukemia virus, MLVenv (18). We therefore hypothesized that the collection of homologous T cell clones expanded following AIP therapy also recognized an epitope from MLVenv. Indeed, the reactivity pattern we observed in our initial screening **(Figure 2A)** closely matched Kang et al.’s finding that B16F10 and MC-38 cells expressed MLVenv while TC-1 cells did not (18).

To further investigate the potential that the dominant anti-tumor response after AIP treatment recognized an endogenous retrovirus (ERV)-derived epitope, we tested several additional cancer cell lines with different levels of ERV expression. It has been shown that ERV transcript numbers can be increased by treating cells with a DNA methyltransferase inhibitor (DNMTi) (19), which we confirmed by treating cells with decitabine and measuring MLVenv transcript level **(Supplementary Figure 2B).** The KP2677 autochthonous lung adenocarcinoma line was found to activate 7PPG2- and 7ELG1-transduced 58^-/-^ T cells with or without DNMTi treatment, while KP-E93-12, 1233 T4, KP-7B, and KP-E85 CC1 W24 cells only activated 7PPG2- and 7ELG1-transduced 58^-/-^ T cells after treatment with decitabine to increase ERV transcript levels **(Supplementary Figure 2C).**

With these data supporting the recognition of an ERV peptide by these T cell clones, we hypothesized that the previously identified MLVenv peptide KSPWFTTL, often referred to as p15E, could be the recognized T cell antigen presented by the class I MHC H-2K^b^ (20). We tested for T cell activation by loading DC2.4 cells with p15E peptide and then coculturing with TCR-transduced 58^-/-^ T cells. SIY peptide was used as a positive control for the 2C T cell line. The 7PPG2- and 7PPG4-transduced 58^-/-^ T cell lines were activated in a dose-dependent manner by the p15E peptide while showing no activation against the SIY peptide even at highest peptide concentration **(Figure 2B).** To further confirm binding, we performed surface plasmon resonance to determine affinity between the pMHCs and several different TCRs. The p15E-K^b^ complex bound TCRs 7PPG2, 7PPG3, 7PPG4, and 7ELG1 with K_D_ values ranging from 20 to 40 μM, within the range of typical affinities observed for TCR-pMHC binding **(Figure 2C)** (21).

### p15E-specific T cells dominate among activated, proliferating cells following AIP therapy

Once we had identified p15E as the antigen recognized by some of the most expanded T cell clones in mice treated with AIP therapy, we further investigated the antigen-specific immune response elicited by AIP treatment. Mice were treated with an updated dosage schedule of the AIP therapy, and p15E-tetramer^+^ CD8^+^ T cells were sorted out from TILs **(Supplementary Figure 3A).** We performed single-cell paired TCR and RNA sequencing of tetramer-positive and tetramer-negative CD8^+^ T cells from AIP-treated mice. In sum, we obtained data for 4,605 tetramer-positive and 3,217 tetramer-negative cells. Strikingly, cells in the tetramer-positive fraction demonstrated substantial levels of clonal expansion relative to cells in the tetramer-negative fraction, indicating that the T cell response to B16F10 tumors undergoing AIP treatment leads to expansion of several clonotypes in response to the p15E antigen **(Figure 3A).**

**Figure 3:**
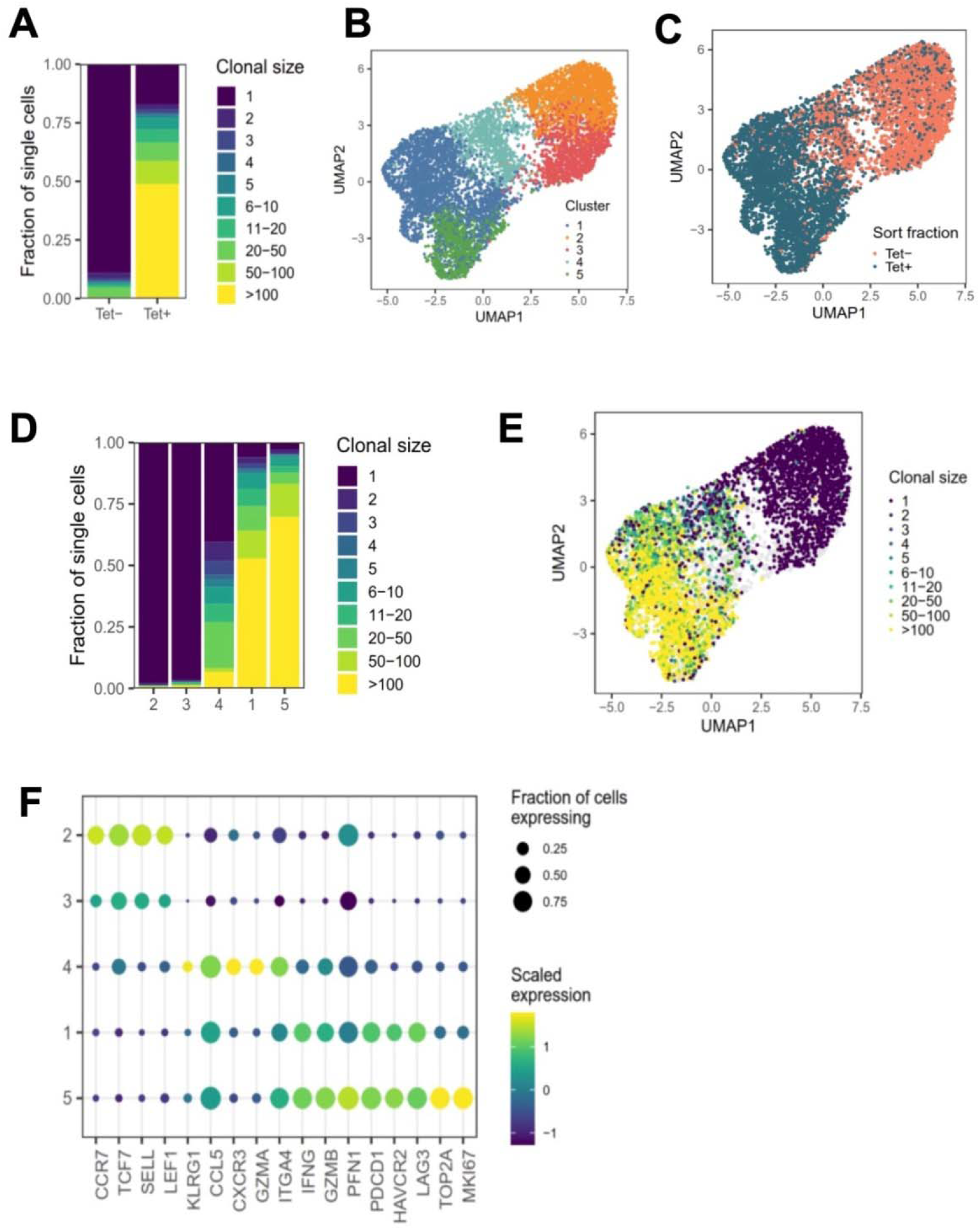
Sequencing of T cells from AIP-treated melanomas reveals significant expansion in the p15E-tetramer^+^ fraction. (A) Clone sizes in tetramer-positive and tetramer-negative populations from TILs isolated from AIP-treated mice as revealed by single-cell T cell sequencing. (B) Unsupervised gene expression analysis of CD8^+^ TILs reveals five distinct transcriptomic clusters. (C) Distribution of tetramer-positive and tetramer-negative cells amongst the 5 clusters. (D) Clusters 1 and 5 show enrichment for expanded clonal TCR sequences. (E) Clone sizes of isolated TILs mapped onto the clusters as determined in B. (F) Expression of a selection of T cell effector and phenotype genes for each transcriptomic cluster. Scaled expression of common genes is indicated by color, and the fraction of cells in that cluster expressing the gene is indicated by relative dot size.

Unsupervised analysis of the gene expression of these cells generated five genotypic clusters (**Figure 3B),** with tetramer-positive cells primarily found in clusters 1, 4, and 5 **(Figure 3C).** Clusters 2 and 3 preferentially expressed *Ccr7* and *Sell* and exhibited little clonal expansion, suggesting that these clusters primarily consist of naïve or bystander T cells **(Figure 3D-F).** Cluster 4 demonstrated an intermediate level of clonal expansion and expressed markers associated with an effector population, including *Klrg1, Cxcr3*, and *Itga4* **(Figure 3D-F).** Both Clusters 1 and 5 were highly clonally expanded and expressed high levels of cytotoxic markers, including *Ifng, Gzmb, Pfn1*, as well as exhaustion markers, such as *Pdcd1, Havcr2*, and *Lag3* **(Figure 3D-F).** In addition to these markers, cluster 5 expressed markers associated with proliferation and cell cycle progression, including *Mki67* and *Top2a* **(Figure 3F).** Cells in the tetramer-positive fraction were 19.7-fold enriched for proliferating cells (Cluster 5) and 21.7-fold enriched for exhausted cells (Cluster 1), indicating that a substantial fraction of CD8^+^ T cells involved in response to B16F10 tumors after AIP treatment are specific for the p15E antigen.

We also analyzed TCR sequences associated with the p15E antigen. In sum, we detected a total of 61 clonally expanded lineages comprised of three or more cells that were at least five-fold enriched in frequency in the tetramer-positive fraction relative to the tetramer negative-fraction. Of these sequences, 37.7% (23 of 61) exhibited usage of the *Traj7* gene segment **(Supplementary Figure 3B).** Among these sequences with *Traj7* usage, we observed a preference for pairing with *Trbv5, Trbv16*, and *Trbv20* **(Supplementary Figure 3C),** and we identified four sequences with a PPG motif in the beta chain and two sequences with a LELGG motif in the beta chain **(Supplementary Figure 3D),** consistent with data obtained from single-cell sorting of T cells. We also identified three sequences with an SWT motif in the beta chain, indicating that other TCR motifs may encode specificity of the p15E epitope **(Supplementary Figure 3D).** In sum, these results demonstrate that the p15E peptide is a dominant T cell antigen in the B16F10 model and that a variety of motifs, in addition to the TRAJ7 alpha chain, can encode specificity for p15E.

### Prophylactic vaccination against p15E does not induce strong T cell responses or delay tumor growth

Given the immunodominance of the p15E antigen following AIP therapy, we next tested whether prophylactic vaccination could delay B16F10 tumor growth. We tested three long peptide vaccine formats with enhanced immunogenicity that have been previously investigated: mouse serum albumin (MSA), transthyretin (TTR), and amphiphile peptide fusions (16,22,23). In each case, a long peptide centered around the p15E 8mer (EGLFNKSPWFTTLISTIMG) was used as the antigenic sequence. C57BL/6 mice were primed with an initial dose of vaccine supplemented by lipo-CpG adjuvant. Mice then received booster vaccines at 14 and 28 days **(Figure 4A).** Intracellular cytokine staining (ICS) of peripheral blood was performed to assess T cell reactivity to p15E at days 21 and 35. The amphiphile vaccine resulted in the highest T cell reactivity, averaging 2.5% among CD8^+^ T cells by day 35 **(Figure 4B).** Although this reactivity was statistically significant, it was modest when compared to amphiphile vaccination with some common model antigens, which can reach upwards of 30% (22,24). Upon tumor challenge with 3×10^5^ B16F10 cells, no delay in tumor growth **(Figure 4C)** or significant increase in survival **(Figure 4D)** was observed.

**Figure 4:**
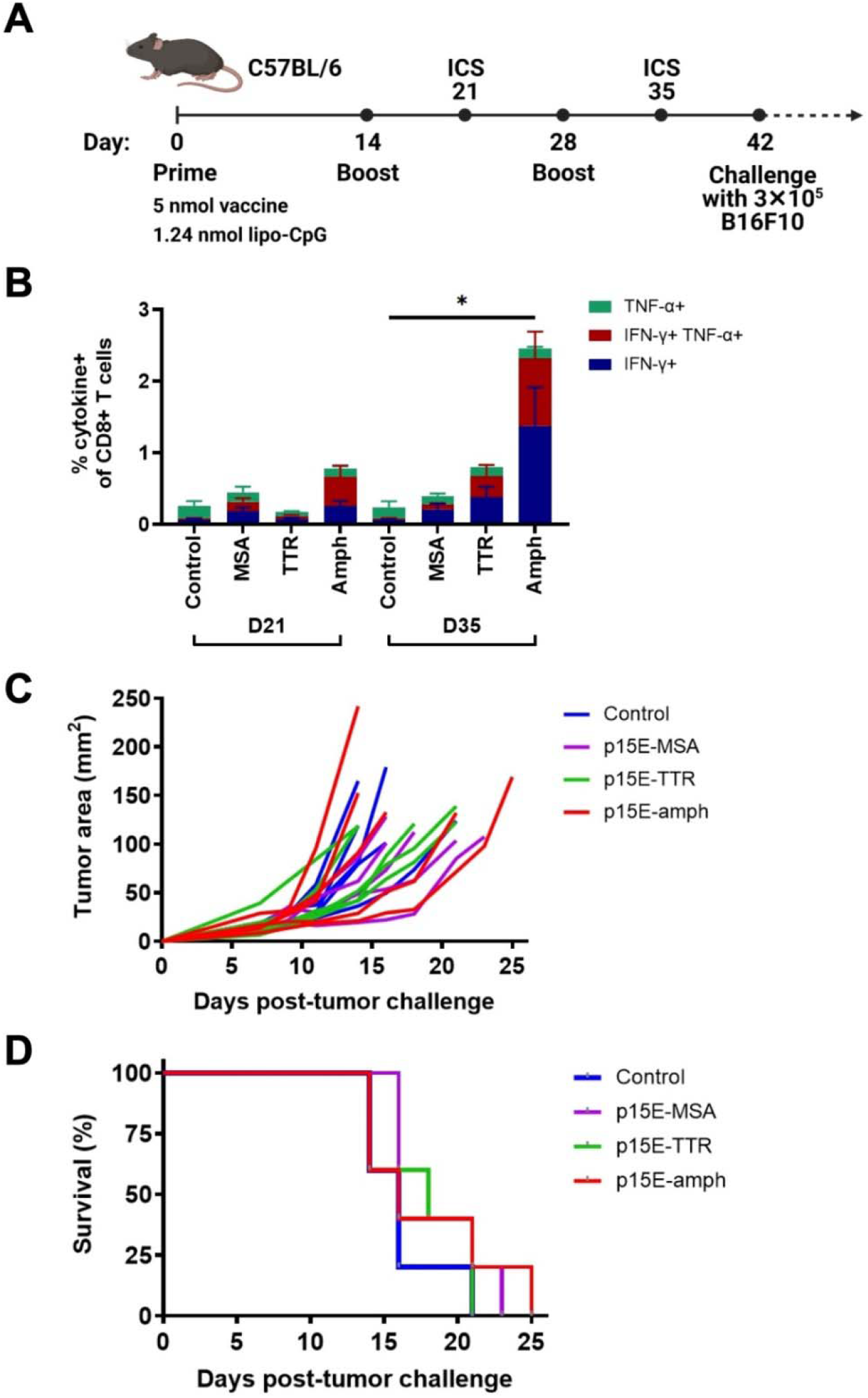
Vaccination against p15E induces minimal expansion of p15E-reactive T cells and does not result in tumor protection. (A) Timeline of vaccination and tumor inoculation. (B) On days 21 and 35, peripheral blood samples from each mouse were stimulated with p15E peptide in the presence of brefeldin A for 4 hours. Intracellular cytokine staining was performed to assess CD8^+^ T cell reactivity to p15E peptide. *P<0.05 by one-way ANOVA with Tukey’s multiple comparisons test. Data shown is mean+s.e.m. (C-D) Tumor areas were measured every other day beginning 3 days post-inoculation. Mice were euthanized when tumor area exceeded 100 mm^2^. Shown are tumor areas for individual mice (C) and survival (D). N=5 mice/group.

### Yeast display of pMHCs allowed identification of mimotopes with increased functionality

We hypothesized that the relatively low T cell response and lack of effect on tumor growth may be due to immune tolerance or thymic deletion caused by low level expression of ERVs in healthy tissues, as has been previously described for self-antigens (25–30). To overcome this limitation, we hypothesized that p15E mimotopes may more potently induce specific T cell activation.

To identify mimotope peptides with increased affinity for the T cell clones, we diversified putative TCR contact residues of the p15E peptide (positions 1, 4, 6, and 7) based upon previously solved H-2K^b^ structures (31,32) and created a library of peptides displayed by H-2K^b^ on yeast **(Supplementary Figure 4A).** Selections were performed on this library using five of the TCRs identified in this study **(Table** 1) and were tracked via next-generation sequencing of enriched yeast following each round of selection. Heat maps were created to show amino acid preference at each position based on read counts of peptides in the sequencing data **(Figure 5A, Supplementary Figure 4B-D).** Generally, position 4 converged to the wild type amino acid tryptophan, with the exception of selection with 7DLG1 which also tolerated tyrosine. Position 1 preferred valine or the wild type lysine, while positions 6 and 7 had more flexibility in amino acid preference depending on the TCR.

**Figure 5:**
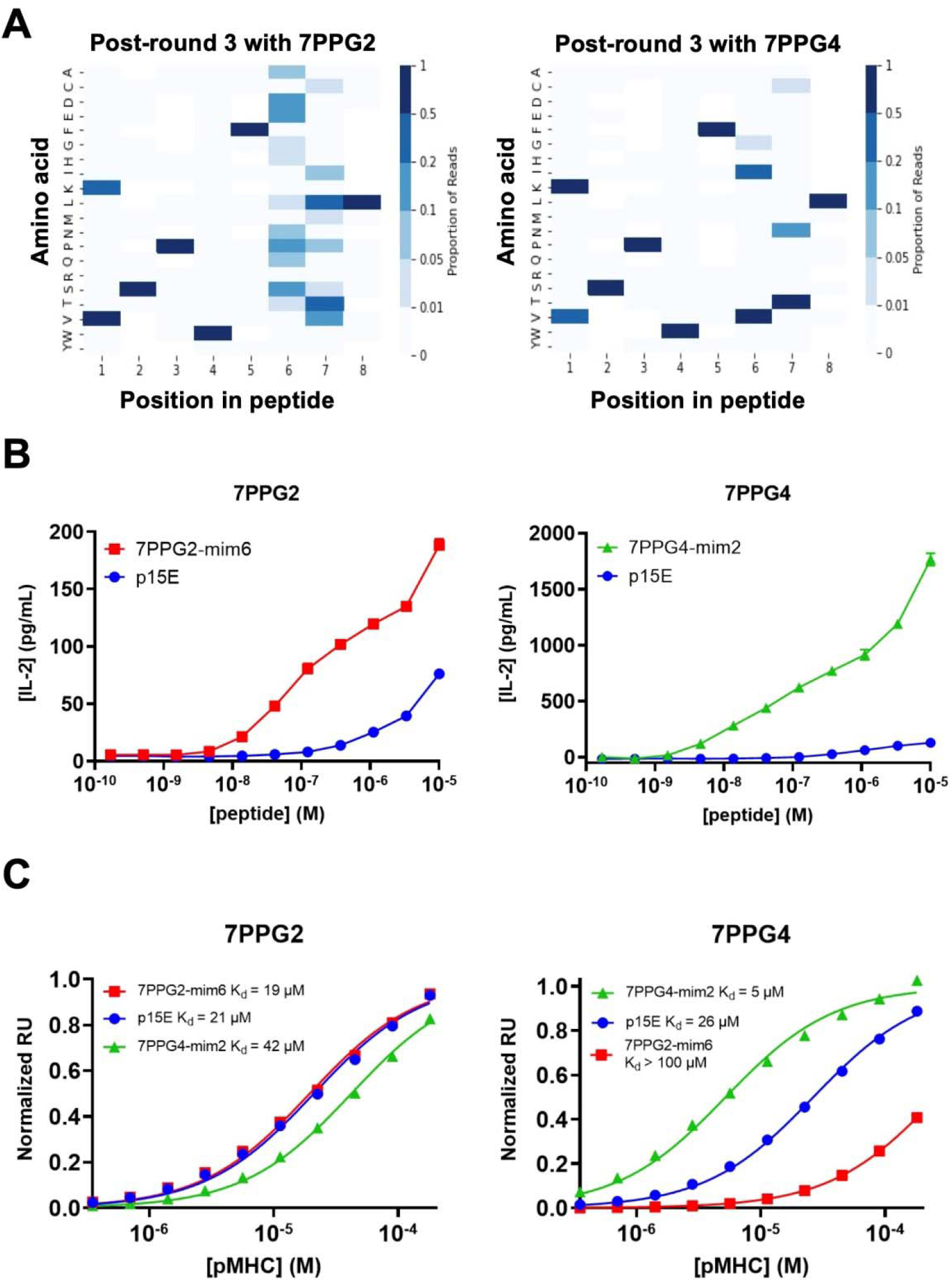
Screening of a yeast displayed peptide-MHC library allowed the identification of mimotopes that more potently activate T cells of interest. (A) Deep sequencing was performed on yeast following each round of selection to reveal amino acid preferences for binding to 7PPG2 and 7PPG4 TCRs. Heat maps show preference for each peptide position, weighted by read count, following 3 rounds of selection with the indicated TCR. (B) TCR-transduced T cells were cocultured with DC2.4 cells loaded with mimotope or p15E peptides. T cell activation was assessed by IL-2 ELISA. Data shown are mean+s.e.m. for triplicate samples and are representative of 3 independent experiments. (C) Surface plasmon resonance was used to quantify affinities between TCRs and mimotope or p15E peptides displayed by H-2K^b^.

We next chose a subset of mimotopes identified from 7PPG2, 7PPG4, and 7ELG1 selections that were either enriched during selections or identified based on the preferences revealed in the heat maps. We tested these mimotopes for their ability to activate their cognate TCR-transduced T cell lines. Several of the mimotopes were able to induce higher IL-2 production in the T cell lines than stimulation with p15E **(Supplementary Figure 5A-C).** The 7PPG2 and 7PPG3 T cells were often more potently activated by peptides with a valine at position 1. 7PPG4 in particular was strongly activated by three mimotopes, with about ten-fold changes over p15E. Based on these results, two mimotopes were chosen for further characterization: 7PPG2-mim6 (VSPWFNTL) and 7PPG4-mim2 (KSPWFITL).

In dose response experiments, each of these mimotopes induced activation much more potently than p15E **(Figure 5B).** To determine if changes to the peptide were affecting pMHC stability, we performed differential scanning fluorimetry on each of the refolded pMHCs. Melting temperatures of the mimotope complexes were within a few degrees of the p15E-K^b^ melting temperature, indicating similar stability **(Supplementary Figure 5D).** We then performed surface plasmon resonance experiments to compare the binding affinities of p15E, 7PPG2-mim6, and 7PPG4-mim2 displayed by H-2K^b^ for TCRs 7PPG2 and 7PPG4. Although 7PPG2-mim6 induced better T cell activation than p15E, the two mimotope peptides had comparable affinities for 7PPG2, ranging from 20 to 40 μM **(Figure 5C).** In contrast, 7PPG4-mim2 had five-fold higher affinity for 7PPG4 than p15E, 5 μM compared to 26 μM **(Figure 5C).** Based on the increased T cell activation induced by both mimotopes and the higher affinity of 7PPG4-mim2, we hypothesized that vaccination with these peptides would induce better T cell responses to p15E and could result in tumor delay.

### Mimotope vaccine-induced T cells can specifically kill p15E-pulsed cells

Following characterization of the mimotope candidates, we performed an *in vivo* killing assay to assess whether vaccination with the mimotopes would result in the expansion of T cells able to kill target cells presenting the p15E antigen. This study consisted of a control group of mice receiving adjuvant only, a p15E amphiphile vaccine group, a group receiving both mimotope vaccines, and an OVA amphiphile vaccine group. Mice received initial vaccines on day 0 and boosts on days 14 and 28 **(Figure 6A).** On day 35, blood samples were taken for ICS to assess antigen-specific T cell expansion, with mice receiving injections of a mixture of dye-labeled peptide-pulsed donor splenocytes **(Figure 6B).** The following day, spleens were removed from experimental mice and processed to assess the remaining populations of dyed cells.

**Figure 6:**
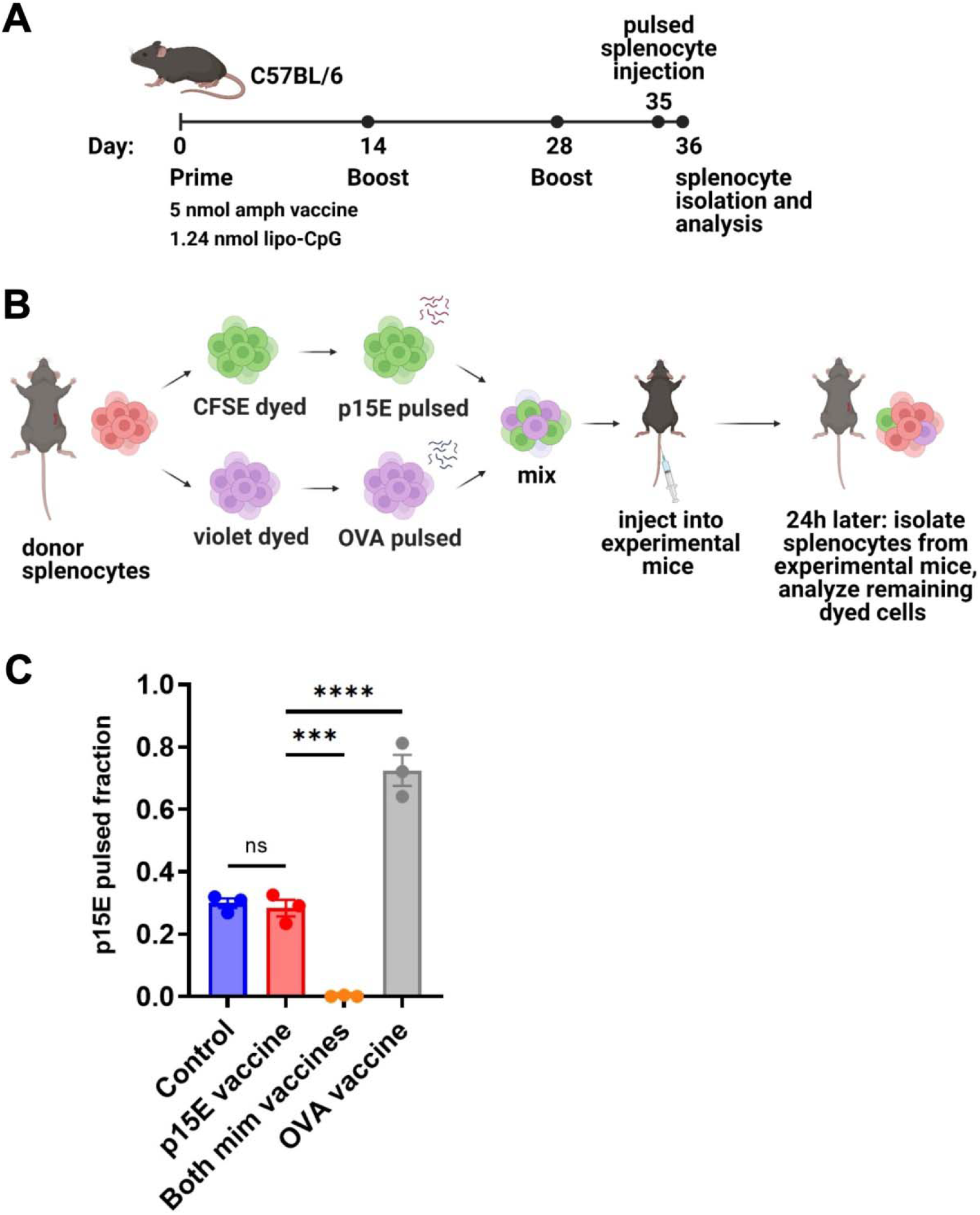
Peptide-pulsed target cells are specifically killed *in vivo* following vaccination with mimotopes. (A) Timeline of vaccination, donor splenocyte injection, and analysis. (B) Schematic of day 35-36 protocol for *in vivo* killing assay. (C) On day 36, spleens were removed from experimental mice. Single cell suspensions were generated and analyzed by flow cytometry for the presence of CFSE- and violet-dyed cells. ns=not significant: P>0.05, ***P<0.001, ****P<0.0001 by one-way ANOVA with Tukey’s multiple comparisons test. Data shown are mean+s.e.m and are representative of 2 independent experiments. N=3 mice/group.

This assay showed that mimotope vaccinated mice were able to specifically kill the p15E-pulsed splenocytes **(Supplementary Figure 6A),** resulting in a very low fraction of p15E-pulsed cells remaining **(Figure 6C).** Similarly, OVA vaccinated mice were able to kill the OVA-pulsed splenocytes, resulting in a high fraction of p15E-pulsed cells remaining. The p15E vaccinated group showed little difference in fraction of p15E-pulsed cells compared to the control group, indicating that no significant killing of either population of peptide-pulsed splenocytes occurred. In terms of reactivity to the targeted antigen, ICS revealed ~5% reactivity to p15E in the group vaccinated with both mimotopes and 20% reactivity to OVA in the OVA vaccinated group **(Supplementary Figure 6B).**

### Vaccination with mimotope peptides increases T cell response to p15E but does not consistently extend survival following tumor challenge

We next sought to assess whether the T cell response induced by mimotope vaccination resulted in tumor protection. For this study, we compared the function of each mimotope individually or the combination of both mimotopes to p15E in a prophylactic vaccination setting. Mice were primed with an initial dose of 5 nmol total amphiphile vaccine (representing 2.5 nmol of each peptide for the group that received both mimotope vaccines) and 1.24 nmol lipo-CpG adjuvant. Booster vaccines were administered at days 14 and 28, with blood drawn for ICS at days 21 and 35 to assess T cell reactivity **(Figure 7A).**

**Figure 7:**
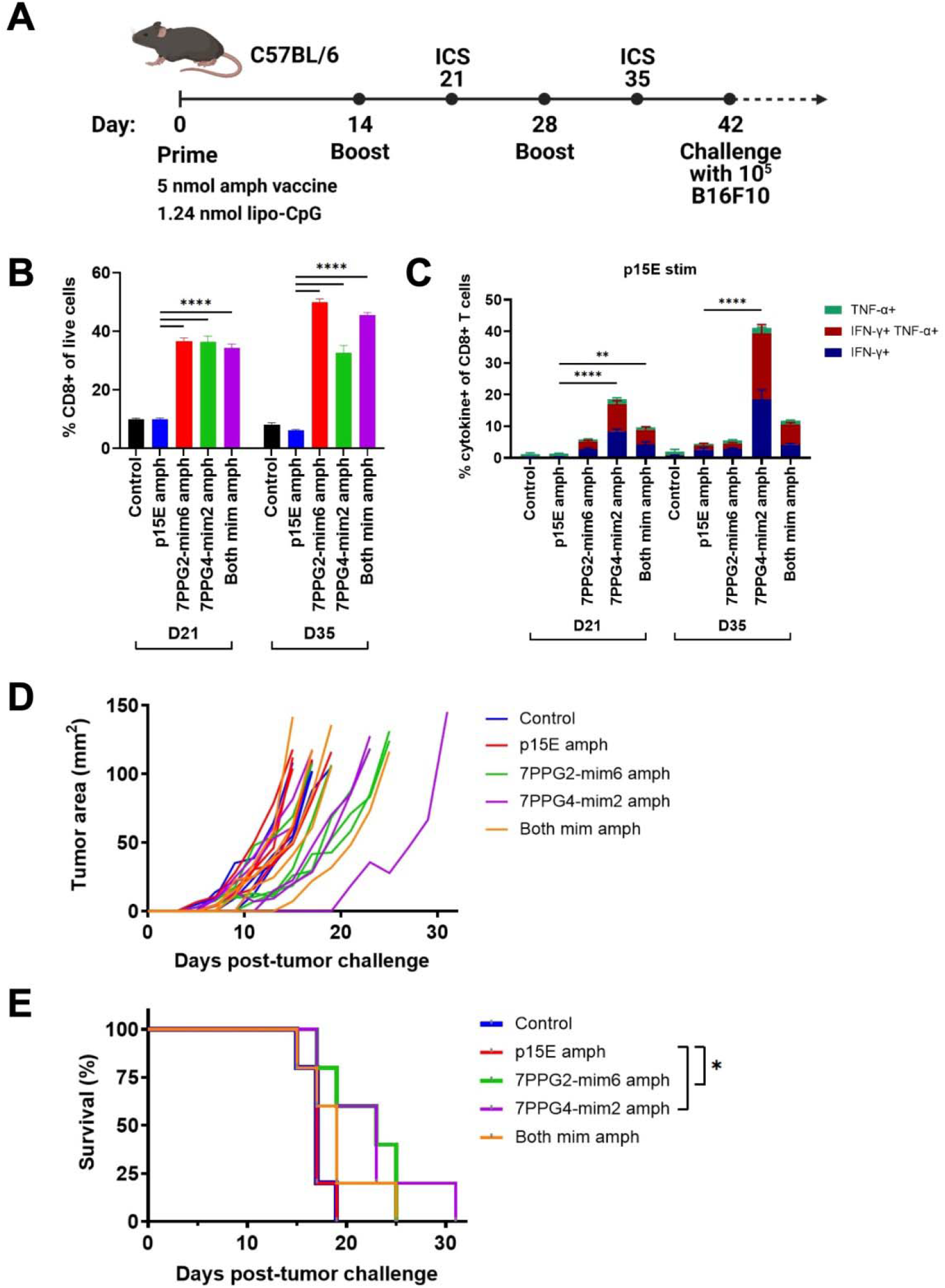
Vaccination against mimotopes promotes the expansion of CD8^+^ T cells reactive to p15E and can result in tumor growth delay. (A) Timeline of vaccination and tumor inoculation. (B-C) On days 21 and 35, peripheral blood samples from each mouse were stimulated with p15E peptide in the presence of brefeldin A for 4 hours. Intracellular cytokine staining was performed to assess the fraction of CD8^+^ T cells within live cells (B) and CD8^+^ T cell reactivity to p15E peptide (C). **P<0.01, ***P<0.001, ****P<0.0001 by one-way ANOVA with Tukey’s multiple comparisons test. Data shown are mean+s.e.m. (D-E) Tumor areas were measured every other day beginning 3 days post-inoculation. Mice were euthanized when tumor area exceeded 100 mm^2^. Shown are tumor areas for individual mice (D) and survival (E). *P<0.05 by log-rank (Mantel-Cox) test. N=5 mice/group.

On day 21, the fraction of CD8^+^ T cells was significantly higher in groups that received any mimotope vaccine as compared to the control or p15E vaccine groups **(Figure 7B).** This disparity increased further by day 35 for 7PPG2-mim6 and both mimotope vaccine groups, indicating that vaccination against the mimotopes was successful at inducing the proliferation of CD8^+^ T cells.

To determine reactivity to specific peptides, blood samples were stimulated with either p15E, 7PPG2-mim6, or 7PPG4-mim2 peptides and assessed for cytokine production via ICS. For mimotope vaccinated groups, there were very high responses to the peptides matched to the vaccine that a group received, ranging from ~25 to 40% of CD8^+^ T cells **(Supplementary Figure 7A, 7B).** Consistent with the increased affinity of 7PPG4 TCR for 7PPG4-mim2 relative to p15E, 7PPG4-mim2 elicited the greatest response, with 40% of CD8^+^ T cells reacting to stimulation at day 21 **(Supplementary Figure 7B).** Mice vaccinated with 7PPG4-mim2 displayed some cross reactivity to 7PPG2-mim6 stimulation, but in contrast, there was minimal reactivity to 7PPG4-mim2 in the group vaccinated with 7PPG2-mim6.

Following stimulation of blood samples with p15E, we found that at both days 21 and 35, the p15E vaccinated group did not show any significant reactivity compared to the control group which received only adjuvant **(Figure 7C).** However, the groups vaccinated with mimotopes displayed higher responses to p15E, in particular the 7PPG4-mim2 vaccinated group which reached 40% reactivity at day 35 **(Figure 7C).** The group vaccinated with both mimotopes generally performed similarly to the 7PPG2-mim6 vaccinated group and worse than the 7PPG4-mim2 vaccinated group when examining reactivity to p15E stimulation.

Six weeks after initial prime **(Figure 7A),** mice were challenged with 1×10^5^ B16F10 melanoma cells on the right flank. Tumors grew at a similar rate in the control and p15E vaccinated groups **(Figure 7D),** resulting in identical survival curves for these two groups **(Figure 7E).** For the individual mimotope vaccine groups, tumors tended to grow more slowly **(Figure 7D),** and these mice had a modest, statistically significant increase in survival **(Figure 7E).** The group vaccinated with both mimotopes contained several mice with delayed tumor growth **(Figure 7D),** although their survival was only modestly different than the p15E vaccinated group **(Figure 7E).** Overall, we did not observe any weight loss over 10% in the period post-tumor challenge **(Supplementary Figure 7C),** indicating that the vaccines were well tolerated. In a second independent experiment, the mimotope vaccinated mice similarly had significantly higher quantities of CD8^+^ T cells and improved reactivity to p15E compared to control and p15E vaccinated groups **(Supplementary Figure 8A-B).** Following tumor challenge, we observed a statistically significant increase in survival for the group receiving both mimotope vaccines, with one mouse completely rejecting the tumor, but no such increase for the two individual mimotope vaccinated groups **(Supplementary Figure 8C).** Thus, we conclude that mimotope vaccination increases the quantity of T cells responsive to p15E but may only provide a modest protective response.

## Discussion

Determining the antigen reactivity of T cells expanded in an anti-tumor immune response can help inform the development of immunotherapies. Thus, we sequenced the TCRs of activated CD8^+^ TILs from B16F10 melanomas and found the T cell response to be oligoclonal in both treated and untreated mice. Other groups have characterized similarly oligoclonal responses for TILs (33,34). Interestingly, there were three TRAV regions represented amongst these clones, but only one J region, TRAJ7. The functionality of TRAJ7 is listed as ORF on IMGT (35), indicating that the gene has an open reading frame (ORF) but has not been functionally described. However, as this region is highly prevalent in our data and we demonstrate proper function of these TCRs, we propose that TRAJ7 is a functional gene. Overall, such similarities in CDR3 sequences are not unheard of; in fact, it has been suggested that public TCRs are prevalent and formed through convergent recombination, especially when there is minimal addition of random nucleotides as observed here (36–39).

We identified the p15E peptide (KSPWFTTL) from the envelope glycoprotein of murine leukemia virus as the antigen recognized by the tumor-reactive T cells expressing homologous TCRs. ERVs arose from retroviral infection of germline cells that resulted in integration into the genome throughout evolution (40). Retrovirus-like elements make up 8-10% of the human and mouse genomes (41). In contrast to the majority of human ERVs, many ERVs in mice remain active (42). C57BL/6 mice carry a single endogenous ecotropic provirus known as *Emv-2* on chromosome 8 (43,44), but it is defective due to a mutation in the reverse transcriptase (45). In healthy tissues, ERV expression is regulated primarily through DNA methylation but also through restriction factors and other epigenetic mechanisms (46–49). In many tumors, however, ERVs are overexpressed (50,51), and *de novo* copies may emerge through recombination of ecotropic and non-ecotropic ERVs, such as melanoma associated retrovirus (MelARV) in B16 tumors (44,52–55).

p15E was previously identified as a T cell antigen (20), with groups demonstrating the expansion of p15E-specific T cells from naive mice and the targeting of p15E in adoptive cell therapy or in vaccine paradigms (20,56–58). A number of previous studies tracking p15E-specific T cells provide evidence for just how dominant p15E can be in anti-tumor immune responses, even in cases in which another antigen is targeted (33,59–61). Our data also support the dominance of the p15E antigen in the case of a combination therapy undirected towards a particular antigen. This recognition appears to utilize highly conserved TCR motifs that can serve as a sequence-based readout of ERV responses. Despite its dominance, our results suggest that even robust p15E-specific responses are not sufficient to induce protective anti-tumor immunity, and it has been previously shown that p15E responses are not necessary for an effective anti-tumor response (61). Tolerance developed against ERVs presents an obstacle to achieving potent anti-tumor immune responses when targeting non-mutated self-antigens such as p15E (62–65), motivating our use of mimotopes to enhance the T cell response.

Characterization of the mimotopes 7PPG2-mim6 and 7PPG4-mim2 revealed their ability to more potently activate their cognate TCRs, but only 7PPG4-mim2 had increased affinity for its TCR compared to p15E. The fact that 7PPG2-mim6 improves potency without increasing affinity for the TCR is notable, but the decoupling of affinity and activity is precedented in previous studies of TCR-antigen systems (10,66–68). While there is previous literature suggesting that increasing the affinity of mimotopes may lead to dysfunction *in vivo* (9), 5 μM is lower than the affinities tested in that case, and 7PPG4-mim2 did not show evidence of deleterious autoreactivity in any *in vivo* studies.

Much of the existing work studying ERVs in mice involves the immunodominant ERV antigen AH1, a peptide from the gp70 surface unit of MLVenv in BALB/c mice (69). AH1 is displayed by the class I MHC H-2L^d^ in CT26 tumors, and it has been successfully targeted in vaccine and adoptive transfer paradigms (57,58,69–71). Additionally, the efficacy of AH1 mimotopes has been well established, with studies demonstrating increased affinities for TCRs, use in prophylactic vaccinations, and cross reactivity within T cell repertoires (8,9,13,14,72–74). Thus, there has been clear demonstration of the benefit of using AH1 mimotopes to improve anti-tumor immunity in BALB/c mice, but to our knowledge, no such identification of p15E mimotopes applicable to C57BL/6 mice. We have demonstrated here the identification of two mimotopes of ERV peptide p15E relevant to multiple tumor types in C57BL/6 mice. The existence of immunodominant responses targeting an ERV epitope in C57BL/6 models treated with immunotherapies such as ICB is particularly notable given how widely used these models are used by cancer biologists and immunologists. Further work is necessary to test these mimotope vaccines in combination with checkpoint inhibitors, MLVenv-targeting antibodies, DNMTis, or other therapeutic modalities, in hopes that they can further improve responses beyond those observed via prophylactic vaccination alone (18,19,75,76).

It should be noted that there are caveats to using mouse models to study ERVs as tumor antigens. While infectious copies of recombined ERVs are common in mice (44,52–55,77), human ERVs (HERVs) so far appear to be inactive (42), which could alter the kinetics of antigen expression. Nevertheless, HERVs have long been studied in association with cancer (78–81). Expression of HERVs has been shown to be a prognostic indicator and to be associated with the response to immunotherapy in renal cell carcinoma and other cancers (82–84). For melanoma specifically, studies have shown elevated HERV-K expression, the production of viral particles, and the presence of HERV-specific antibodies in patient sera (85–87). Further, Schiavetti et al. reported the identification of an HERV-K antigen presented by HLA-A2 and recognized by CD8^+^ T cells in melanoma patients (88).

There is substantial evidence that HERVs can be therapeutically targeted as TAAs in some cancers. Chimeric antigen receptor (CAR) T cells targeting HERVs have been developed for breast cancer and melanoma (89,90), and adoptive transfer of HERV-E specific T cells is being evaluated in a phase I clinical trial for renal cell carcinoma (91). Finally, in a study particularly relevant to the translational potential of ERV-targeting vaccines, Sacha et al. vaccinated rhesus macaques with simian ERV-K Gag and Env and found that vaccination induced T cell responses with no adverse effects or induction of autoimmune disease (92). These studies, in conjunction with our results, suggest that both natural and engineered anti-ERV immune responses present potential novel avenues towards anti-cancer immunotherapies.

## Materials and Methods

### Mice

4-6 week old female C57BL/6 mice were obtained from the Jackson Laboratory (C57BL/6J) or Taconic (C57BL/6NTac) and used in studies beginning at 6-8 weeks of age. All animal work was conducted under the approval of the MIT Division of Comparative Medicine in accordance with federal, state, and local guidelines.

### Cells and media

HEK293-T cells were obtained from ATCC. B16F10 and DC2.4 cells were gifts from the Irvine lab. B16.SIY cells were a gift from the Spranger lab. MC-38 and TC-1 cells were gifts from the Wittrup lab. KP lines were gifts from the Jacks lab. CD8^+^ 58^-/-^, Sf9, and Hi5 cells were gifts from the Garcia lab.

HEK 293-T, B16F10, B16.SIY, and MC-38 cells were cultured in Dulbecco’s Modified Eagle’s Medium (ATCC) supplemented with 10% fetal bovine serum (FBS, R&D Systems), 100 U/mL penicillin (Thermo), and 100 μg/mL streptomycin (Thermo). TC-1, DC2.4, and 58^-^/^-^ lines were cultured in RPMI-1640 media (ATCC) supplemented with 10% FBS, 100 U/mL penicillin, and 100 μg/mL streptomycin. All mammalian cell lines and assay cultures were maintained at 37°C and 5% CO_2_. All cells were frequently tested and confirmed negative for mycoplasma contamination. B16F10 cells used for tumor challenge studies also tested negative for rodent pathogens.

Sf9 cells were cultured in Sf-900 III SFM (Gibco) supplemented with L-glutamine, 10% FBS (R&D Systems), and 20 μg/mL gentamicin sulfate (Lonza). Hi5 cells were cultured in Insect-XPRESS (Lonza) supplemented with 10 μg/mL gentamycin sulfate (Lonza). Insect cell lines were maintained at 27°C with shaking at 120 rpm.

### Flow cytometry

Antibodies to Myc (clone 9B11) and Flag (clone D6W5B) were purchased from Cell Signaling Technology. Antibodies to human β_2_M (clone 2M2), mouse TCR β chain (clone H57-597), mouse H-2K^b^/H-2D^b^ (clone 5041.16.1), mouse IFN-γ (clone XMG1.2), mouse TNF-α (clone MP6-XT22), mouse CD8α (clone 53-6.7), and mouse CD16/32 (Fc block, clone 93) were purchased from Biolegend. Other reagents used include Zombie Aqua Viability Dye (Biolegend), CellTrace CFSE and CellTrace Violet (Invitrogen). Samples were run on an Accuri C6 (BD), LSRII (BD), or Cytoflex (Beckman Coulter) and analyzed in the Accuri software or FlowJo v10. Samples were run in FACS buffer (phosphate-buffered saline (PBS) with 2 mM EDTA and 0.5% bovine serum albumin (BSA)).

### T cell sorting and sequencing following AIP therapy

Mice were inoculated with 10^6^ B16F10 cells subcutaneously on the right flank. On days 8 and 15, mice received AIP therapy (TA99, extended half-life IL-2, and anti-PD-1) as previously described (16). On day 21, mice were euthanized, single cell suspensions were generated from tumors, and red blood cells were lysed. IFN-γ secreting cells were labeled using the mouse IFN-γ secretion assay (Miltenyi Biotec) and then stained for viability and CD8. Splenocytes stimulated with 10 ng/mL PMA and 10 μg/mL ionomycin were used as a positive control. IFN-γ^+^ CD8^+^ T cells were sorted into RT-PCR buffer in 96 well plates, one cell per well, on a FACSAria (BD).

Single-cell sequencing of TCRs was performed by nested reverse-transcription polymerase chain reaction (RT-PCR) using previously published primer sets (93,94). RT-PCR was performed using One Step RT-PCR kits (Qiagen) and a mixture of TRAV-, TRBV-, TRAC-, and TRBC-specific primers. Nested PCR was then performed with a second set of TCR alpha- and beta-specific primers. A third PCR step was then performed separately for alpha and beta chains to add group, plate, column, and row DNA barcodes, as well as P5 and P7 sequences for paired end NGS. Alpha and beta chain PCR products were then pooled separately, run on 1% agarose gels, and purified by gel extraction (GeneJET PCR Gel Extraction Kit, Thermo). Samples were sequenced at the MIT BioMicroCenter using Illumina MiSeq 500nt paired-end kits.

TCR sequencing data was analyzed using a combination of in-house and publicly available programs. Contigs were assumed from paired end reads by Flash (95) and an in-house program was used to demultiplex sequences and segregate alpha and beta chain sequences. Unique sequences were then collapsed using MIGEC (96) and sequences were fed through HighV-Quest (97) to assess VDJ usage and CDR3 sequence. Paired chain sequences were assembled based on demultiplexed read barcodes. TCRs in separate wells with identical CDR3 sequences and VDJ usages were identified as members of a clonal lineage. Any contaminating sequences from previous sequencing runs were manually identified through comparison of exact nucleotide sequences and examination of sequences which appeared in wells that intentionally did not contain T cells.

### p15E-specific T cell sorting, sequencing, and phenotyping

Mice were inoculated with 10^6^ B16F10 cells subcutaneously on the right flank. At day 8, mice received AIP therapy, followed by ICB every three days thereafter, as previously described for 1× AIP therapy (98). On day 26, mice were euthanized. Single cell suspensions were generated from tumors and spleens. Pan-T cell isolations were performed using an EasySep mouse T cell isolation kit (Stem Cell), and then T cells from three tumors were pooled and stained with Zombie Aqua, Fc block, 5 nM p15E-tetramer (produced in-house), CD3, and CD8. Splenocytes were used as controls for sorting.

### Single-cell RNA sequencing

Tetramer-positive and tetramer-negative CD8^+^ T cells were processed for single-cell RNA sequencing using the Seq-Well platform with second strand chemistry (99,100). Libraries were barcoded and amplified using the Nextera XT kit (Illumina) and were sequenced on the Illumina Novaseq.

### Analysis of single-cell gene expression

Raw read processing was performed as in Macosko et al (101). Briefly, sequencing reads were aligned to the mm10 reference genome, collapsed by unique molecular identifier (UMI) and aggregated by cell barcode to obtain a gene expression matrix of cells versus genes. Barcodes with under 300 unique genes detected were then excluded from analysis. The data was then log-normalized by library size and variable genes were selected using the FindVariableGenes function in Seurat. The ScaleData function in Seurat was then used to scale the data to unit variance using a Poisson model and to regress out the number of genes detected per cell (102). The RunPCA function was then used to perform principal component analysis, and the top 20 principal components were used to generate a UMAP with the RunUMAP function. Clusters were then determined with the FindClusters function.

### Recovery and analysis of TCR sequences from single cell libraries

TCR sequences were recovered from TCR libraries according to Tu et al. (103). Briefly, biotinylated baits for TRBC and TRAC were used to enrich TCR transcripts from barcoded whole-transcriptome amplification product. This product was then further amplified using mouse V-region primers and prepared for sequencing using Nextera sequencing handles. Libraries were sequenced on an Illumina Miseq using 150 nt length reads. CDR3 sequences were obtained as described previously.

### Peptides

Crude peptides used for T cell activation experiments and pMHC refolds were obtained from the Proteomics Core of the Swanson Biotechnology Center. Peptide sequences are as follows: p15E: KSPWFTTL, 7PPG2-mim6: VSPWFNTL, 7PPG4-mim2: KSPWFITL, SIY: SIYRYYGL, OVA: SIINFEKL.

### T cell activation assays

#### T cell line generation

For creation of T cell lines, TCR alpha chain was linked to TCR beta chain by a P2A sequence, followed by EGFP fluorescent protein sequence, in the MP71 retroviral vector. 80% confluent HEK293-T cells were transfected with 2 μg of TCR plasmid and 1 μg pCL-Eco helper plasmid with 6 μg (3× DNA mass) PEI (Santa Cruz Biotech) to create retrovirus. 2 days later, the supernatant was then used to transduce CD8^+^ 58^-^/^-^ cells with 0.8 μg/mL polybrene (Santa Cruz Biotech). Cells were spinfected at 1000×g and 32°C for 1.5 hours. 2 days later, transduction was assessed by expression of EGFP and by staining with an antibody against murine TCR β chain. If necessary, cells were sorted on EGFP on a FACSAria (BD) to obtain a >95% transduced population.

#### Tumor cell co-culture

Recombinant murine IFN-γ (Peprotech) was added to a working concentration ≥500 U/mL (1:1000 from a 0.1 mg/mL stock solution) to tumor cell lines 24 hours before co-culture setup to increase MHC expression. 25,000 T cells and tumor cells at an E:T ratio of 1:1, 2:1, 5:1, or 10:1 per well were co-cultured in 200 μL RPMI complete media in 96 well plates.

#### DC co-culture

25,000 DC2.4 cells per well were incubated with various concentrations of peptide in 100 μL RPMI complete media for 2-3 hours before the addition of T cells. 25,000 T cells per well were added in 100 μL RPMI complete media in 96 well plates.

After 24 hours, plates were spun down at 500×g for 5 minutes, and 2×100 μL samples of supernatant were saved and frozen at −80°C. T cell activation was assessed using mouse IL-2 ELISA kits (Invitrogen), with samples diluted 1:2 −1:20 for comparison to the standard curve. Absorbance values were measured with a Tecan Infinite m200 Pro plate reader.

### RT-qPCR

Total RNA was collected from cultured cells treated with 100 nM 5-aza-2⍰-deoxycytidine (decitabine; Sigma) or splenocytes from a naïve C57BL/6 mouse using NucleoSpin RNA Plus (Takara). RNA was reverse transcribed with SuperScript III RT and random primers (Invitrogen) to synthesize first-strand cDNA. RT-qPCR was performed with Luna Universal qPCR reagent (NEB). eMLV env was detected with 5’-AGGCTGTTCCAGAGATTGTG-3’ and 5’-TTCTGGACCACCACATGAC-3’ and 18S rRNA was detected with 5’-GTAACCCGTTGAACCCCATT-3’ and 5’-CCATCCAATCGGTAGTAGCG-3’ on Roche LightCycler 480 II (104,105).

### TCR expression and purification

Recombinant soluble TCRs used for yeast library selections and surface plasmon resonance experiments were produced using a baculovirus expression system in Hi5 insect cells. TCR alpha chain extracellular domain sequences were cloned into pAcGP67a vector containing a 3C protease site (LEVLFQGP) followed by an acidic leucine zipper, AviTag biotinylation site (GLNDIFEAQKIEWHE), and poly-histidine tag (6×His). TCR beta chain extracellular domain sequences were cloned into pAcGP67a vector containing a 3C site followed by a basic leucine zipper and poly-histidine tag. 2 μg of each plasmid was transfected into SF9 insect cells with BestBac 2.0 linearized baculovirus DNA (Expression Systems) using Cellfectin II reagent (Invitrogen). SF9s were then infected with 1:1000 primary transfection supernatant to amplify the virus. Titration of alpha and beta chain viruses was performed to identify the optimal ratio of the two viruses for heterodimer formation. Protein was then produced by large scale infection of Hi5 cells. Secreted protein was purified using HisPur Ni-NTA resin (Thermo) for immobilized metal affinity chromatography (IMAC) and biotinylated overnight at 4°C using biotin (Avidity), ATP, and BirA ligase. Proteins were further purified by size exclusion chromatography via an S200 column on an AKTApure FPLC (Cytiva). Biotinylation efficiency was assessed by adding streptavidin (ThermoFisher) to ~2 ug of protein and running an SDS-PAGE gel. Aliquots of biotinylated TCR were stored in HBS with 20% glycerol at −80°C.

### pMHC protein expression and purification

The following were separately cloned into bacterial expression vector pet28a: the extracellular domains α_1_, α_2_, and α_3_ of heavy chain H-2K^b^ plus an AviTag biotinylation site and poly-histidine tag, and human β2M plus a poly-histidine tag. Inclusion bodies for H-2K^b^ and β2M were generated in BL21 *E. coli* and denatured in 8 M urea and 6 M guanidine hydrocholoride. Excess peptide and β2M were injected into MHC refolding buffer (100 mM Tris, 400 mM L-arginine hydrochloride, 2 mM EDTA, 0.5 mM oxidized glutathione, 5.0 mM reduced glutathione, 0.2 mM PMSF). H-2K^b^ was injected one hour later at a 1:1 mass ratio with β2M, and the solution was incubated for 24 hours at 4°C. Complexes were desalted by dialysis in deionized water at room temperature over 2 days and then purified using IMAC or anion exchange followed by size exclusion chromatography on an AKTApure FPLC (Cytiva). For SPR experiments, complexes were purified using an S200 column followed by an S75 column and used fresh. For pMHC to be used in formation of pMHC tetramers, complexes were biotinylated overnight at 4°C using biotin (Avidity), ATP, and BirA ligase and then purified using an S200 column; monomers were stored in HBS at −80°C. pMHC tetramers were created by adding pMHC monomer to tetrameric AlexaFluor647-conjugated streptavidin (produced in-house as previously described) at a 6:1 ratio and incubating for 30 minutes at 4°C (106).

### Surface plasmon resonance

Steady-state surface plasmon resonance experiments were performed with a Biacore T200 instrument. TCRs were immobilized at 1100 – 2400 response units on Series S CM5 sensor chips (Cytiva) by amine coupling. pMHC was injected as analyte with a concentration range of 0.35 – 266 μM and a flow rate of 10 μL/min at 25°C. Samples were in HBS-EP (0.01 M HEPES pH 7.4, 0.15 M NaCl, 3 mM EDTA, 0.005% v/v Surfactant P20) or HBS-EP+ buffer (0.01 M HEPES pH 7.4, 0.15 M NaCl, 3 mM EDTA, 0.05% v/v Surfactant P20). Data was fit with GraphPad Prism 9 one site specific binding model and normalized to R_max_.

### Differential scanning fluorimetry

Differential scanning fluorimetry (DSF) was performed using a CFX384 RT-PCR instrument (Bio-Rad). Excitation and emission wavelengths were 587 and 607 nm, respectively. Samples were at 10 μM pMHC concentration with 10× SYPRO Orange (Invitrogen) in 10 μL HBS-EP+ buffer in a 386 well plate. The temperature range was 20 to 95°C with a scan rate of 1°C per minute. Measurements were performed on triplicate samples.

### Vaccine studies

MSA and TTR long peptide fusions were produced in house by the Wittrup lab as previously described (23). Long peptide-amphiphile fusion vaccines were obtained from Northwestern University Peptide Synthesis Core (p15E and mimotopes) or produced in house by the Irvine lab as previously described (OVA amph) (16,22). Lipo-CpG was produced in house by the Irvine lab as previously described (16,22). 5 nmol of vaccine and 1.24 nmol of lipo-CpG per injection were used in a volume of 100 μL PBS, sterile filtered, and injected subcutaneously at the tail base, 50 μL per side, using 31G insulin syringes. Mice were primed and then boosted at 2 and 4 weeks following prime (n=5 per group).

Vaccine sequences are as follows:

MSA-p15E: MSA-EGLFNKSPWFTTLISTIMG
TTR-p15E: TTR-EGLFNKSPWFTTLISTIMG
p15E amph: DSPE-PEG2000-CEGLFNKSPWFTTLISTIMG
7PPG2-mim6 amph: DSPE-PEG2000-CEGLFNVSPWFNTLISTIMG
7PPG4-mim2 amph: DSPE-PEG2000-CEGLFNKSPWFITLISTIMG

### Intracellular cytokine staining (ICS)

One week following each vaccine boost, blood samples were drawn retro-orbitally from each mouse. 50-80 μL was used per well for stimulation with peptide, and excess blood from each group was pooled and used for a positive and negative control well per group. Red blood cells were lysed with ACK lysis buffer (Thermo), and then cells were washed once with FACS buffer. In 100 μL/well RPMI complete media, sample wells were stimulated with 10 μg/mL peptide (p15E, 7PPG2-mim6, or 7PPG4-mim2), and positive control wells were stimulated with 10 ng/mL PMA and 10 μg/mL ionomycin. Cells were stimulated for two hours before 1 uL/well brefeldin A (BD GolgiPlug) was added, and then cells were incubated for an additional four hours. Cells were washed twice with cold PBS and then stained with Zombie Aqua 1:1000 and Fc block 1:100 in 100 μL/well PBS for 15 minutes at room temperature. Cells were then stained with CD8 APC 1:100 in 100 μL/well FACS buffer for 20 minutes at room temperature. Cells were fixed and permeabilized using 100 μL/well of CytoFix/CytoPerm solution (BD) for 20 minutes at 4°C. Following two washes with Perm/Wash buffer (BD), cells were stained with IFN-γ PE and TNF-α AF488 1:75 each in 50 μL/well Perm/Wash buffer for 30 minutes at 4°C. Following two washes with FACS buffer, samples were run on an LSR II (BD) or Cytoflex (Beckman Coulter) and data analyzed in FlowJo v10. Outliers were identified using the ROUT method with Q= 1% (GraphPad Prism 9).

### Tumor challenge

Two weeks after the second vaccine boost, 10^5^ B16F10 cells in 100 μL PBS were injected subcutaneously on the right flank using 29G insulin syringes. Tumor areas were calculated by multiplying length and width taken by caliper measurement every other day beginning three days post-tumor inoculation. Mice were euthanized when tumor area exceeded 100 mm^2^.

### In vivo killing assay

Donor mice were euthanized and spleens removed. In a sterile hood, single cell suspensions were created by mashing spleens through 70 μm mesh filters. Red blood cells were lysed using ACK lysis buffer (Thermo), and cells were washed twice with PBS. Cells were dyed with either CellTrace CFSE (Invitrogen) for 8 minutes at 2.5 μM or CellTrace Violet (Invitrogen) for 20 minutes at 5 μM, quenched with 10% FBS, and washed twice with lymphocyte media (RPMI-1640 with HEPES and L-glutamine, 10% FBS, 1% pen-strep, 1× NEAA, 1 mM sodium pyruvate, and 50 μM β-mercaptoethanol). Cell concentration was adjusted to 5×10^6^ cells/mL, and 10 μg/mL of either p15E or OVA peptide was added. Cells were incubated for two hours at 37°C in 5% CO_2_. Cells were washed twice with cold PBS and mixed roughly 1:1 CFSE-dyed (p15E pulsed) to violet-dyed (OVA pulsed) cells. Around 5×10^6^ cells in 100 μL were injected per experimental mouse via the tail vein using 29G insulin syringes (n=3 per group). 24 hours later, experimental mice were euthanized and splenocytes analyzed to compare ratios of CFSE-dyed to Violet-dyed cells. Samples were run on a Cytoflex and data analyzed in FlowJo v10.

### Expression of H-2K^b^ on the surface of yeast

Yeast displayed H-2K^b^ was designed as a single chain trimer cloned into the pYAL vector. The Aga2 signal peptide directly precedes the peptide, which is linked to human β2M linked to heavy chain extracellular domains α_1_, α_2_, and α_3_ linked to Aga2. Flexible glycine-serine linkers were used as follows: 3×GGGGS between peptide and β2M, 4×GGGGS between β2M and MHCI heavy chain, and a Myc epitope tag (EQKLISEEDL) followed by 3×GGGGS between MHCI heavy chain and Aga2. The binding groove of the MHC is opened via a tyrosine to alanine mutation at position 84 of the heavy chain (Y84A) to accommodate the linker connecting the peptide to β2M (107). The Myc tag at the C-terminal end of the pMHC construct allows antibody labeling and detection by flow cytometry. In addition, the Aga2p protein fused to the C-terminus of the pMHC construct allows association with Aga1p on the surface of yeast via disulfide bridges (108).

Plasmid was transformed into EBY100 yeast by electroporation. Yeast were grown in SDCAA media pH 4.5 for 1 day at 30°C, shaking at 250 rpm, and induced in SGCAA media pH 4.5 for 2 days at 20°C, shaking at 250 rpm (10,109). Construct expression was assessed by flow cytometry following staining with a Myc tag antibody.

Proper folding of the construct was verified using a tag enrichment experiment, in which yeast displaying p15E-K^b^ with a Myc tag were mixed with Flag-tagged yeast at a 1:10 ratio. Yeast were stained with Myc and Flag antibodies to verify that the Myc-tagged yeast were present at about 10%. Streptavidin beads (Miltenyi Biotec) were loaded with 400 nM 7PPG2 TCR and then incubated with the yeast mixture. The solution was selected using an LS column (Miltenyi Biotec). A sample of selected yeast was run on a flow cytometer to verify that the selected population was majority p15E-K^b^ yeast expressing the Myc tag, indicating proper folding of the pMHC and successful binding to the TCR.

### Mimotope library design and selection

To create a mimotope library, potential TCR contact residues of the p15E peptide (KSPWFTTL) were randomized. The library was designed as XSPXFXXL, where X was a position randomized using an NNK codon during polymerase chain reaction (PCR) on the MHC construct. The starting H-2K^b^ template did not contain a peptide so as to avoid contaminating the library with a template peptide sequence. For creation of a library of at least 10^8^ transformants, RJY100 yeast were electroporated with a 5:1 mass ratio mixture of PCR product and linearized pYAL vector. Yeast were grown for 1 day in SDCAA media at 30°C, shaking at 250 rpm, and then induced for 2 days in SGCAA media at 20°C, shaking at 250 rpm. The library was selected for 3 rounds with streptavidin beads (Miltenyi Biotec) loaded with biotinylated TCR and then for 1-2 additional rounds with tetramers of TCR and anti-AlexaFluor647 microbeads (Miltenyi Biotec). TCR tetramers were created by adding TCR to tetrameric AlexaFluor647-conjugated streptavidin (produced in-house) at a 5:1 ratio and incubating for 10 minutes at 4°C.

### Yeast deep sequencing and analysis

DNA was obtained from at least 10^7^ yeast per round of selection using a Zymoprep Yeast Plasmid Miniprep II kit (Zymo Research). Primers were designed to amplify the Aga2 leader peptide through to the beginning of the β2M sequence, covering the peptide encoding region, by PCR. A second round of PCR served to add the i5 and i7 anchors for Illumina sequencing as well as a barcode unique to a given round of selection. Amplicons were sequenced by the MIT BioMicroCenter on an Illumina MiSeq using a 300nt v2 kit for 150nt paired-end reads. Paired-end reads were assembled using PANDASeq (110), clustered using CD-HIT (111), and further analyzed for amino acid prevalence at each position (weighted by read count) using an in-house program.

### Statistical analysis

GraphPad Prism 9 software was used for all statistical analyses. Data with replicates is shown as the mean value with error bars showing standard error of the mean (s.e.m.). Tests, p-values, and group/replicate sizes are indicated in figure captions.

## Data Availability

The original TCR sequencing and mimotope library selection datasets generated for this study can be found in NCBI’s Sequence Read Archive under BioProject ID PRJNA754131 (https://www.ncbi.nlm.nih.gov/sra/PRJNA754131). The followup p15E-specific TCR sequencing and phenotyping data discussed in this publication have been deposited in NCBI’s Gene Expression Omnibus and are accessible through GEO Series accession number GSE196782 (https://www.ncbi.nlm.nih.gov/geo/query/acc.cgi?acc=GSE196782).

## Conflicts of Interest

B.E.G. is currently an employee of Repertoire Immune Medicines. M.E.B. is a co-founder, equity holder, and advisor of Viralogic Therapeutics and Abata Therapeutics, an equity holder in 3T Biosciences, and an inventor on patents related to yeast display approaches licensed to 3T Biosciences. D.J.I. is a co-founder, SAB member, and equity holder in Elicio Therapeutics, and is an inventor on patents licensed to Elicio Therapeutics related to the amphiphile-vaccine technology. J.C.L. has interests in Honeycomb Biotechnologies, and is an inventor on patents related to the SeqWell technology. M.E.B., D.J.I, and J.C.L’s interests are reviewed and managed under MIT’s policies for potential conflicts of interest. The remaining authors declare that the research was conducted in the absence of any commercial or financial relationships that could be construed as a potential conflict of interest.

## Author Contributions

Study conception: B.E.G. and M.E.B. Experimental design and execution: B.E.G., C.M.B., D.M.M., B.H.K., N.K.S., B.D.H., C.G.R., K.D.M., L.M., C.S.D., T.K., K.S.G., O.C.T.M., D.G., S.W. Data analysis: B.E.G., D.M.M, N.K.S., B.D.H., C.G.R., P.V.H. Supervising research: J.C.L., K.D.W., D.J.I., M.E.B. Writing manuscript: B.E.G. and M.E.B. Editing manuscript: all authors.

## Funding

This work was funded by the Melanoma Research Alliance (to K.D.W. and M.E.B.). This work was supported in part by the Koch Institute Support (core) Grant P30-CA14051 from the National Cancer Institute. M.E.B. was additionally supported by funding from the Packard Foundation, Schmidt Futures, the Pew Foundation, the V Foundation, AACR-TESARO Career Development Award for Immuno-oncology Research [17-20-47-BIRN]. B.E.G. was additionally supported by the National Institutes of Health Biotechnology Training Program (5T32-GM008334), the National Science Foundation Graduate Research Fellowship, and the Margaret A. Cunningham Immune Mechanisms in Cancer Research Fellowship Award. C.M.B. was supported by a postdoctoral fellowship from the Ludwig Center at MIT’s Koch Institute for Integrative Cancer Research.

## Acknowledgements

Parts of some figures were created with BioRender.com. We thank Naveen Mehta and Christine Devlin for their assistance in designing experimental reagents and the Spranger, Jacks, and Garcia labs for providing cell lines and protocols. We thank Dr. Megan Burger for helpful discussion. We would like to thank the Koch Institute’s Robert A. Swanson (1969) Biotechnology Center for their technical support, especially the Flow Cytometry Facility, MIT BioMicro Center, and Biopolymers and Proteomics Core. We also thank the Division of Comparative Medicine at MIT. We would like to thank Mark Karver and the Peptide Synthesis Core Facility at Northwestern University for production of amphiphile vaccine reagents. This facility is currently supported by the Soft and Hybrid Nanotechnology Experimental (SHyNE) Resource (NSF ECCS-2025633) and has also been supported by the Simpson Querrey Institute for BioNanotechnology, Northwestern University Office for Research, U.S. Army Research Office, and the U.S. Army Medical Research and Materiel Command.

