## Supplemental Data for "Identification of highly cross-reactive mimotopes for a public T cell response in murine melanoma"

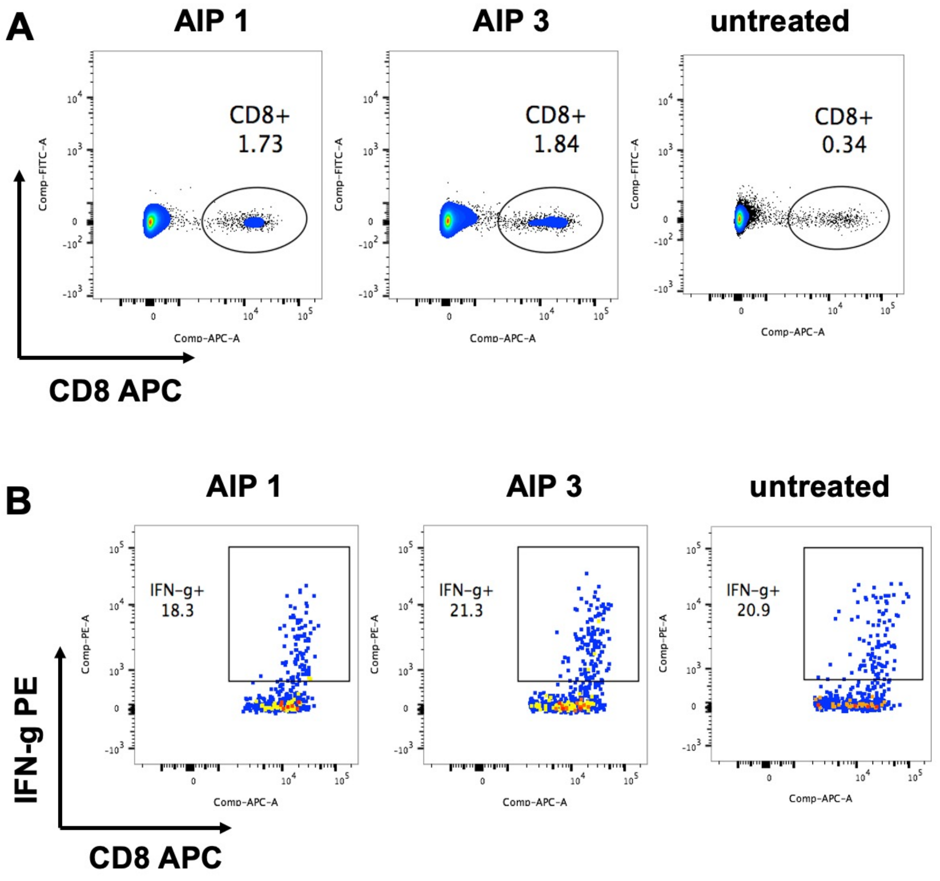


C

| TRAV | CDR1 | CDR2 |
|---|---|---|
| 5D-4 | NSASNY | IRSNMER |
| 7-2 | DRNVDY | IFSNGE |
| 10/10D | DTASSY | IRSNVDR |

**Supplementary Figure 1:**
(A-B) Stained single-cell suspensions of B16F10 tumors were assessed for CD8$^{+}$ T cell fraction (A) and IFN-γ$^{+}$ fraction (B). (C) CDR1 and CDR2 sequences for the three variable regions found in the set of similar T cell clones from B16F10 melanomas.

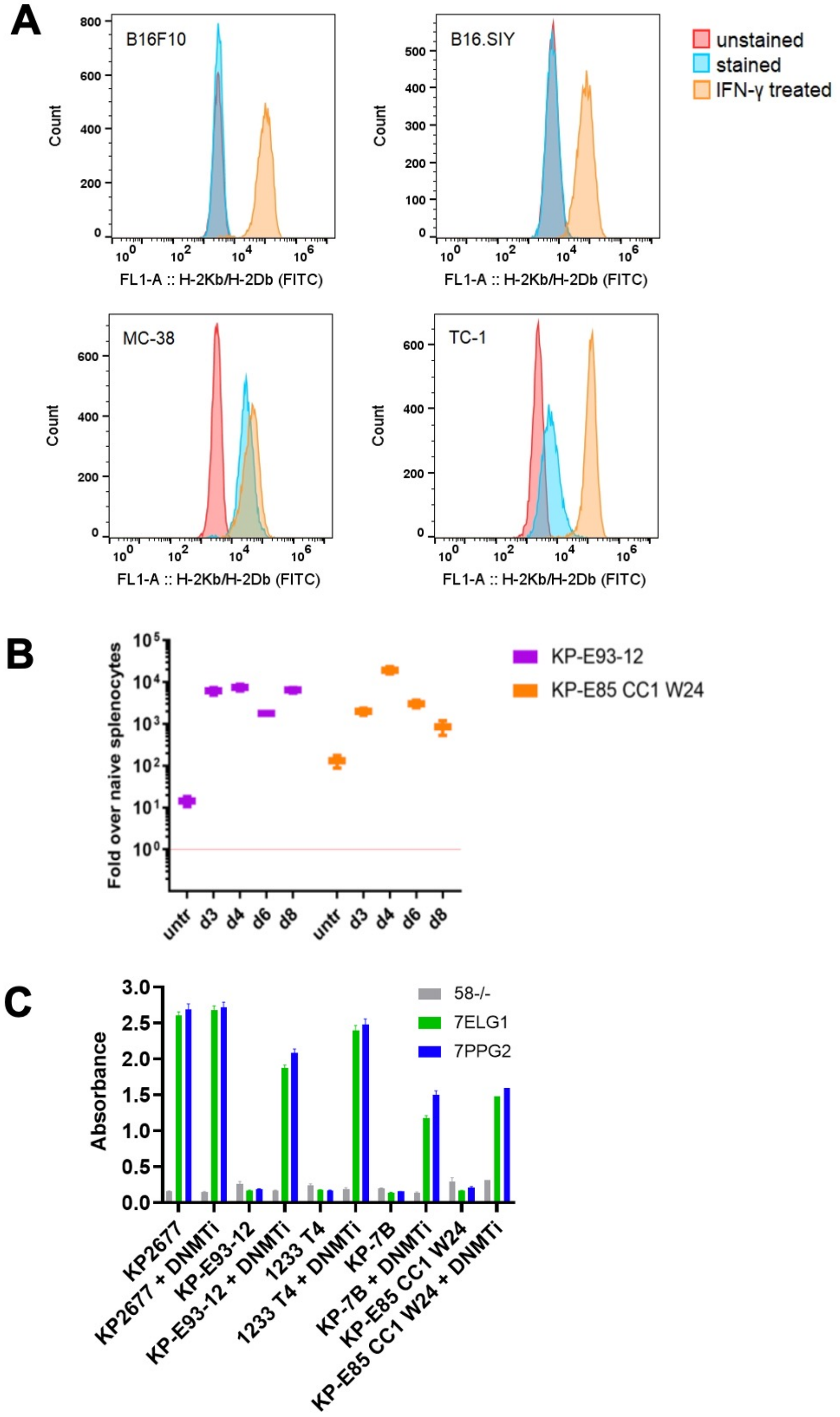


**Supplementary Figure 2:**
(A) Staining of cancer cells lines to assess MHC class I expression before and after 24 hour treatment with IFN-γ. (B) Quantification of MLVenv transcript levels following DNMTi treatment. untr=untreated, d=day. (C) Untransduced or TCR-transduced $58^{-/-}$ cells were cocultured with cancer cell lines after treatment with DNMTi for 4 days. T cell activation was assessed by IL-2 ELISA. Data shown are mean+s.e.m. for duplicate ($58^{-/-}$) or triplicate samples (7ELG1, 7PPG2), except for KP-E85 CC1 W24 + DNMTi which are single data points.

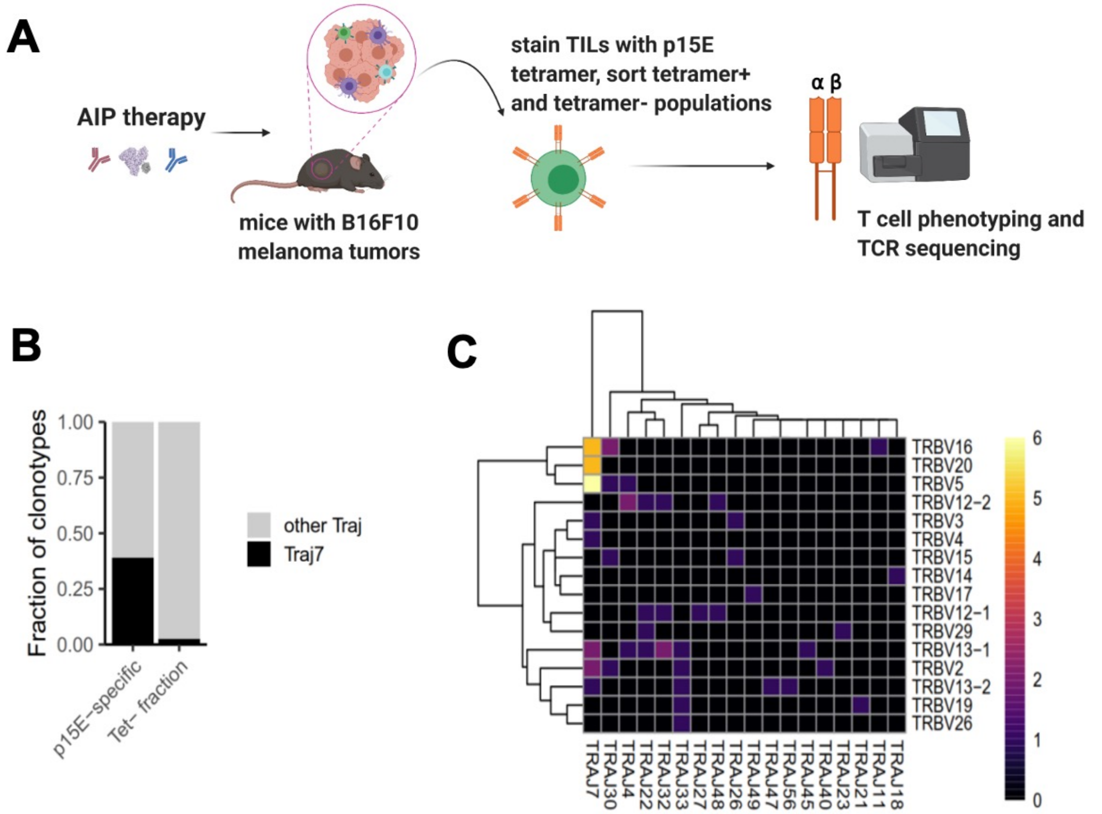


**D**

| Clonal Size | CDR3α | TRAV | TRAJ | CDR3β | TRBV | TRBJ |
|---|---|---|---|---|---|---|
| 57 | CAAKDYSNNRLT | 7-2 | 7 | CASS**PPG**GGDEQYF | 5 | 2-7 |
| 24 | CALSEDYSNNRLT | 12-2 | 7 | CASS**PPG**GGTEVFF | 5 | 1-1 |
| 16 | CAAQDYSNNRLT | 7-2 | 7 | CASS**PPG**AGTGQLYF | 5 | 2-2 |
| 8 | CAMREAGAFNYGNEKITF | 16D/DV11 | 48 | CASS**PPG**GWGQNTLYF | 12-1 | 2-4 |
| 8 | CATDEDYSNNRLT | 8-1 | 7 | CASS**LELGG**LEQYF | 16 | 2-7 |
| 6 | CAVTPDYSNNRLT | 3-1 | 7 | CASS**LELGG**QEQYF | 16 | 2-7 |
| 10 | CALGEGSNYQLIW | 6-6 | 33 | CAS**SWT**DSDYTF | 26 | 1-2 |
| 7 | CAVRGNYAQGLTF | 3D-3 | 26 | CAS**SWT**GTTSAETLYF | 3 | 2-3 |
| 5 | CAASGDTNAYKVIF | 14D-3/DV8 | 30 | CAS**SWT**GGDTQYF | 15 | 2-5 |

**Supplementary Figure 3:**
(A) Schematic showing process to acquire T cells and perform sequencing and phenotyping. (B) Fraction of clonotypes using *Traj7* versus other variable regions. (C) Pairing preferences for TRAJ and TRBV regions. (D) TCR sequences for T cell clones specific for p15E and expanded following AIP therapy of B16F10 melanomas.

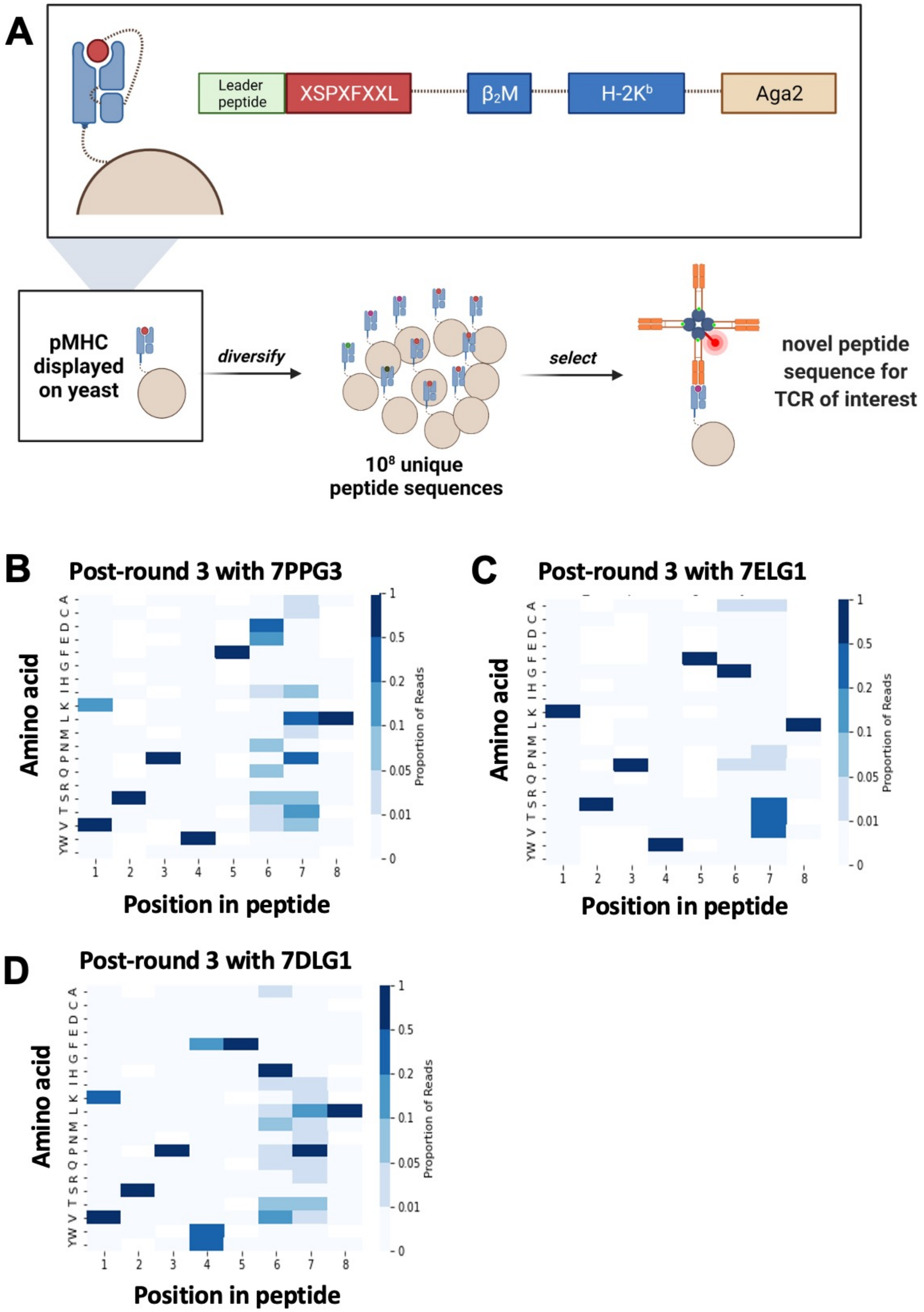


**Supplementary Figure 4**:
(A) Schematic of peptide-MHC yeast display library creation and selections with TCR. (B-D) Deep sequencing was performed on yeast following each round of selection. Heat maps show the amino acid preference at each peptide position, weighted by read count, following three rounds of selection with the indicated TCR.

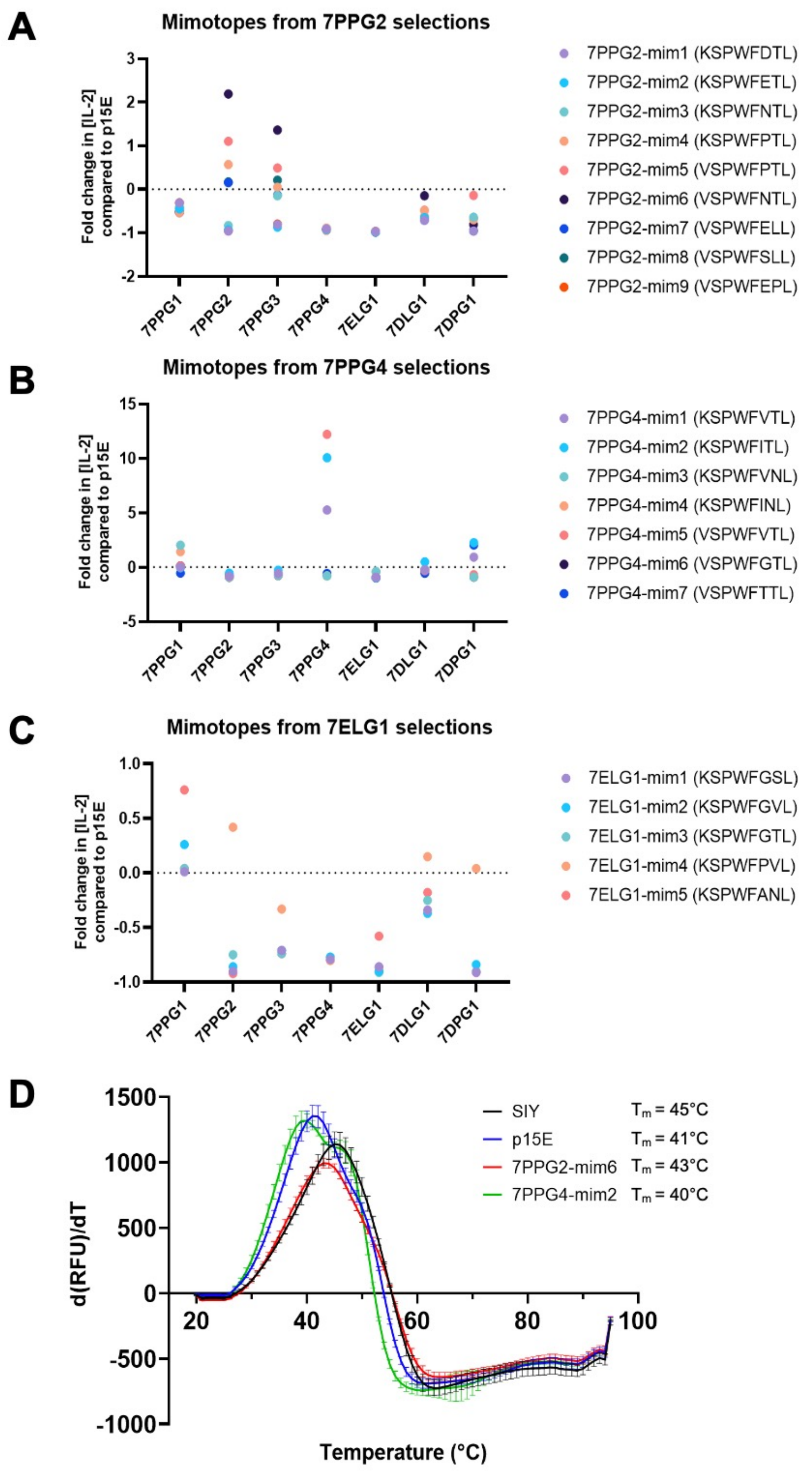


**Supplementary Figure 5:**
(A-C) TCR-transduced 58$^{-/-}$ cells were cocultured with DC2.4 cells loaded with mimotope or p15E peptide. T cell activation was assessed by IL-2 ELISA. Fold change in IL-2 concentration was calculated as [mimotope/p15E]-1. (D) Differential scanning fluorimetry was used to compare stability of H-2K$^b$ in complex with the mimotope peptides, p15E, and SIY. Data shown is mean+s.e.m. of triplicate samples.

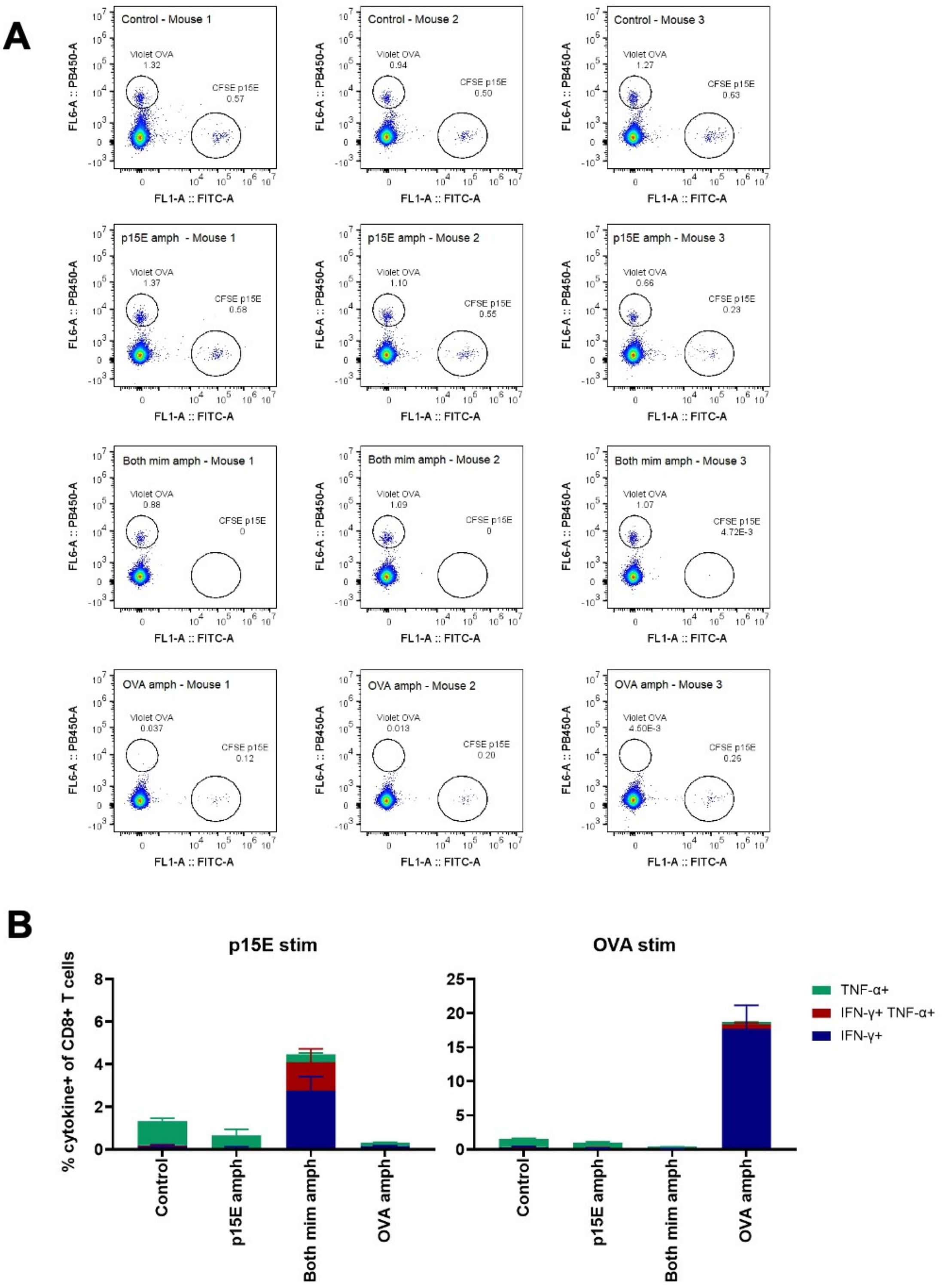


**Supplementary Figure 6:**

(A) Flow cytometry plots showing dyed cell populations. (B) On day 35, peripheral blood samples from each mouse were stimulated with p15E or OVA peptide in the presence of brefeldin A for 4 hours. Intracellular cytokine staining was performed to assess CD8+ T cell reactivity to each peptide. Data shown are mean+s.e.m. N=3 mice/group.

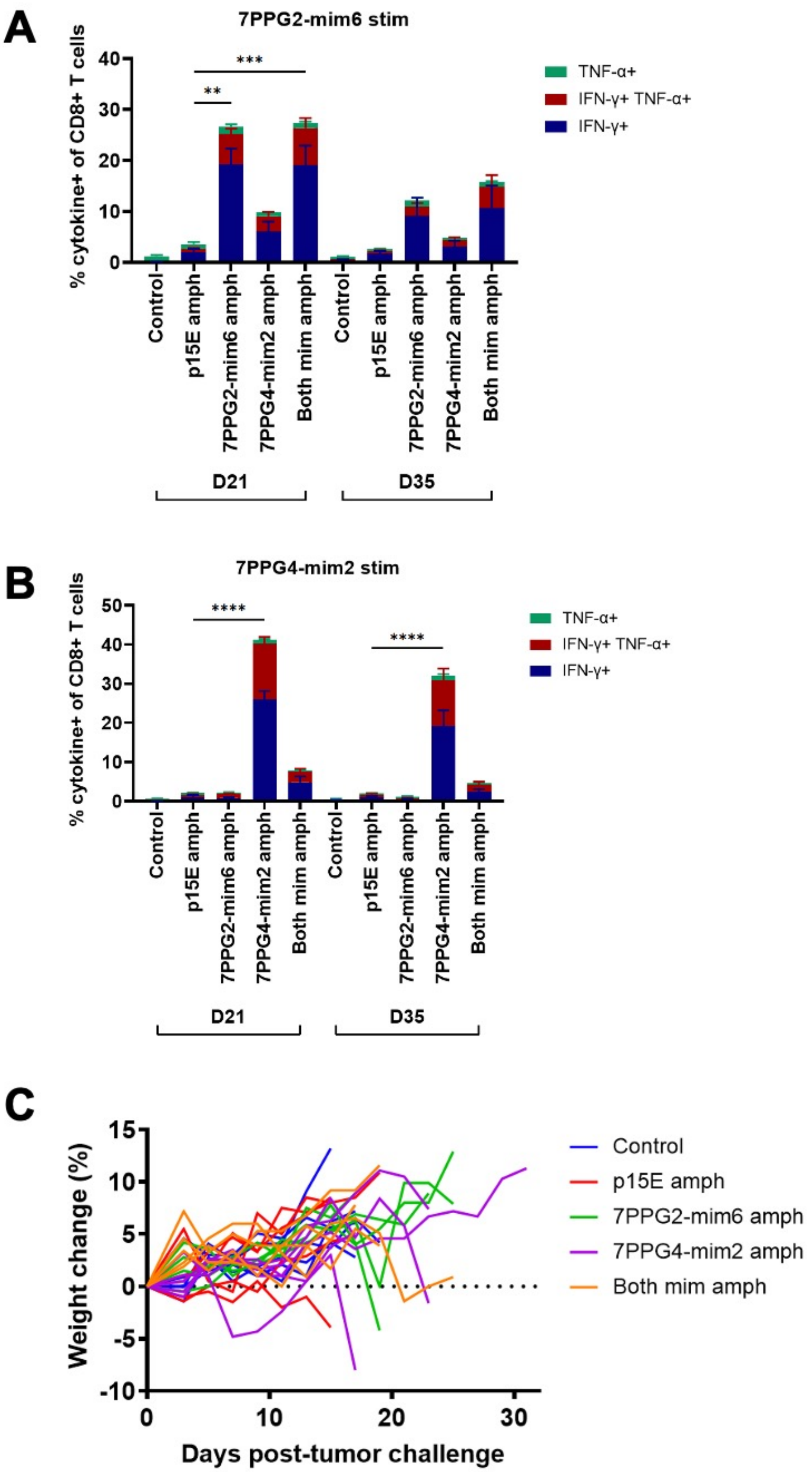


**Supplementary Figure 7:**

(A-B) On days 21 and 35, peripheral blood samples from each mouse were stimulated with the indicated mimotope peptide in the presence of brefeldin A for 4 hours. Intracellular cytokine staining was performed to assess CD8$^{+}$ T cell reactivity to each peptide. **P<0.01, ***P<0.001, ****P<0.0001 by one-way ANOVA with Tukey's multiple comparisons test. Data shown are mean+s.e.m. (C) Weight change as a percent of starting weight for individual mice following tumor inoculation. Weight was measured every other day beginning three days post-tumor inoculation. N=5 mice/group.

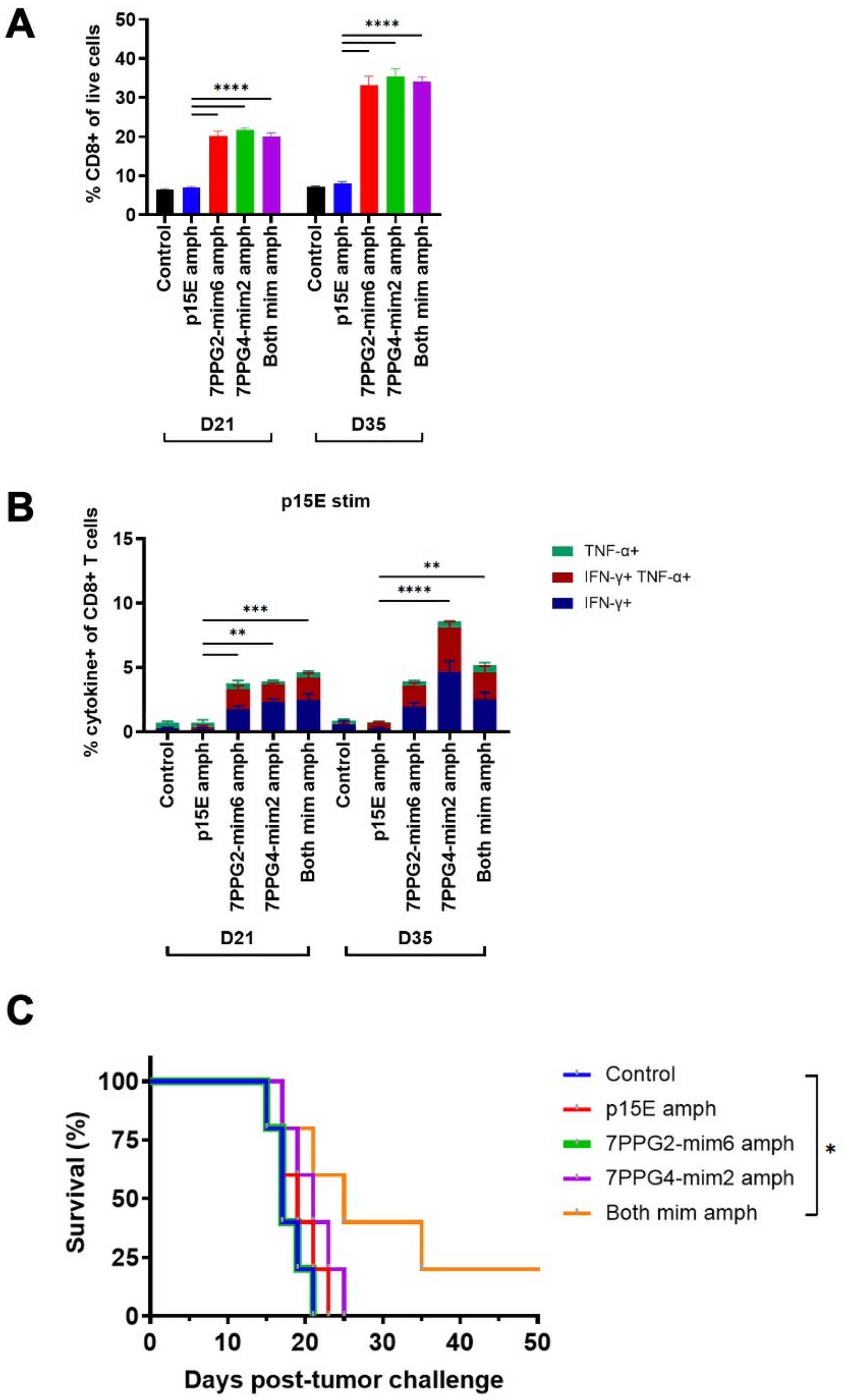


**Supplementary Figure 8:**
(A-B) On days 21 and 35, peripheral blood samples from each mouse were stimulated with p15E peptide in the presence of brefeldin A for 4 hours. Intracellular cytokine staining was performed to assess the fraction of CD8+ T cells within live cells (A) and $CD8^+$ T cell reactivity to p15E peptide (B). **P<0.01, ***P<0.001, ****P<0.0001 by one-way ANOVA with Tukey's multiple comparisons test. Data shown are mean+s.e.m. (C) In a second independent study, tumor areas were measured every other day beginning three days post-inoculation. Shown are survival curves. Mice were euthanized when tumor area exceeded 100 $mm^2$. *P<0.05 by log-rank (Mantel-Cox) test. N=5 mice/group.
